# AnnoPRO: an Innovative Strategy for Protein Function Annotation based on Image-like Protein Representation and Multimodal Deep Learning

**DOI:** 10.1101/2023.05.13.540619

**Authors:** Lingyan Zheng, Shuiyang Shi, Pan Fang, Hongning Zhang, Ziqi Pan, Shijie Huang, Weiqi Xia, Honglin Li, Zhenyu Zeng, Shun Zhang, Yuzong Chen, Mingkun Lu, Zhaorong Li, Feng Zhu

## Abstract

Protein function annotation has been one of the longstanding issues, which is key for discovering drug targets and understanding physiological or pathological process. A variety of computational methods have therefore been constructed to facilitate the research developments in this particular direction. However, the annotation of protein function based on computational methods has been suffering from the serious “*long-tail problem*”, and it remains extremely challenging for existing methods to improve the prediction accuracies for protein families in *tail label levels*. In this study, an innovative strategy, entitled ‘*AnnoPRO*’, for protein function annotation was thus constructed. **First**, a novel method enabling image-like protein representations was proposed. This method is unique in capturing the intrinsic correlations among protein features, which can greatly favor the application of the *state-of-the-art* deep learning methods popular in image classification. **Second**, a multimodal framework integrating multichannel convolutional neural network and long short-term memory neural network was constructed to realize a deep learning-based protein functional annotation. Since this framework was inspired by a reputable method used in image classification for dealing with its ‘*long-tail problem*’, our *AnnoPRO* was expected to significantly improve the annotation performance of the protein families in *tail label level*. Multiple case studies based on benchmark were also conducted, which confirmed the superior performance of *AnnoPRO* among the existing methods. All source codes and models of *AnnoPRO* were freely available to all users at https://github.com/idrblab/AnnoPRO, and would be essential complement to existing methods.

## 1. Introduction

Protein function annotation has been one of the longstanding issues, which is key for discovering new drug target and understanding physiological or pathological process [1-3]. With the advance of next-generation sequencing, a large amount of protein sequences have been accumulated, and over 200 million sequences have been available in UniProt [4]. Compared with protein sequences, the acquirement of experimentally-validated protein functions is much more challenging, which is characterized by its nature of time-consuming & labor-intensive [5-7]. So far, the functions of only 90 thousand proteins have been successfully annotated, which asks for the development of new strategies to significantly accelerate the process of protein function annotations [8-10]. Thus, a variety of computational methods have been constructed to facilitate the research developments in this particular direction [11-14], which successfully promote the identification of efficacy drug target, the revealing of molecular mechanism underlying disease etiology, and so on [15-18].

However, the annotation of protein function using computational method has been suffering from the serious “*long-tail problem*” [15-18]. As shown in **Fig. 1**, the total number (5,323) of *Gene Ontology* (GO) families in the ‘*Tail Label Levels*’ is more than 10 times larger than that (459) of the ‘*Head Label Levels*’ [19]. In other words, the protein function data in GO database follow a *long-tailed distribution* where only a few ‘*head label*’ families and many ‘*tail label*’ ones exist [19]. The ‘*long-tailed phenomenon*’ has been reported to lead to severe degradation of annotation performances due to the serious imbalance problem between the data of *head* and *tail* [20]. This is also the principal reason for *head label* families dominating the training process, making these families enjoy much higher accuracies than the *tail label* ones [19-21]. To cope with this problem, two types of protein annotation strategy have been proposed, which can be roughly divided into the sequence homology (SH) based and the machine learning (ML) based ones [22-24].

**Fig. 1.**
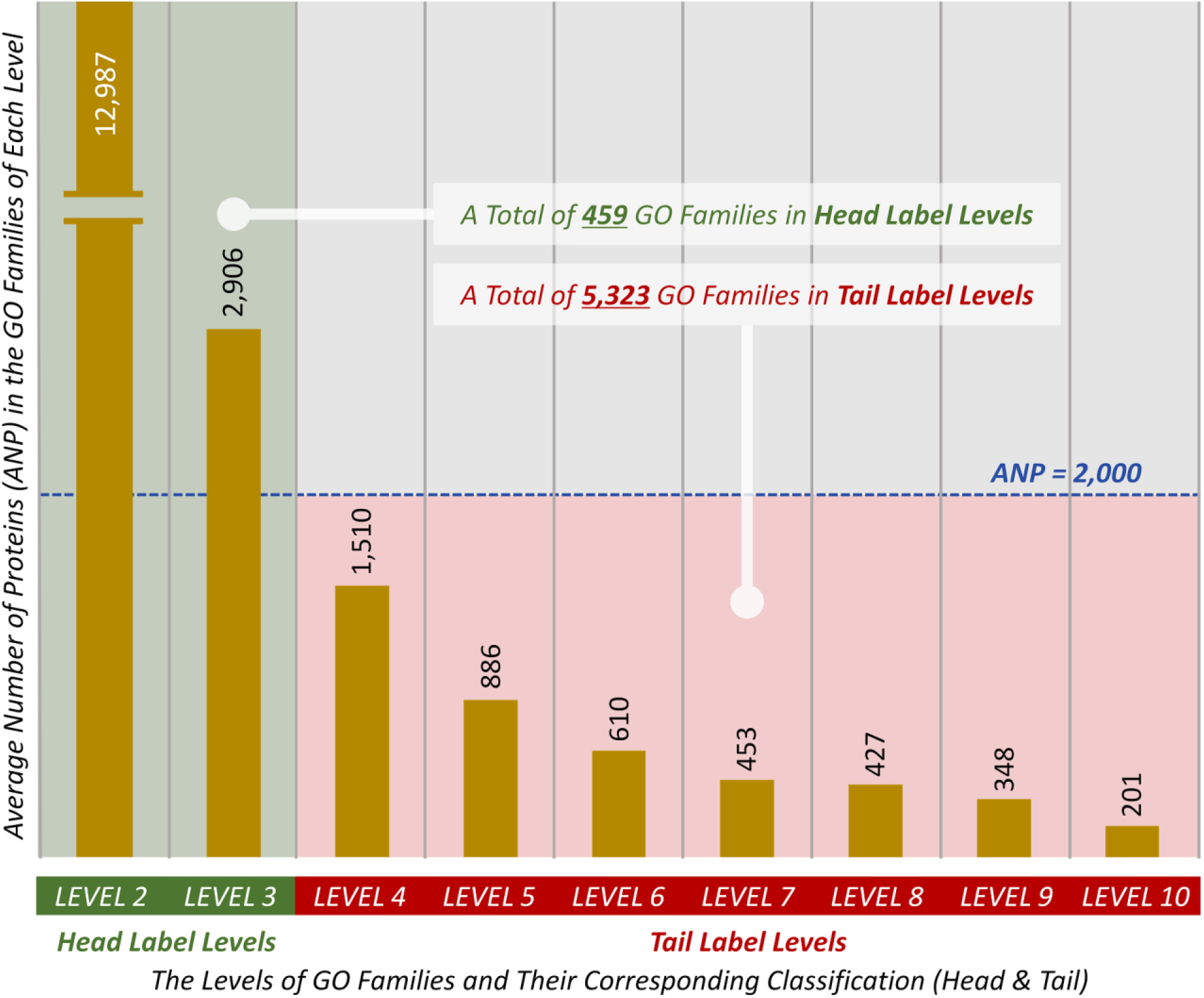
Average numbers of proteins (ANP) in the GO families of nine different levels (LEVEL 2 to LEVEL 10 as shown in **Supplementary Fig. S2**). There was a clear descending trend of ANPs from the top level (LEVEL 2) to the bottom one (LEVEL 10). Since the ANP of one family indicated its representativeness among all families, this figure denoted a gradual decrease of the representativeness of a family with the penetration into deeper level. Therefore, these nine levels could be classified into two groups based on their ANPs: the “*Head Label Levels*” (ANP of their GO families ≥ 2,000) and the “*Tail Label Levels*” (ANP of their GO families < 2,000). As shown, the total number (5,323) of GO families in the “*Tail Label Levels*” was >10 times larger than that (459) of the “*Head Label Levels*”, and such kind of data distribution induced a serious ‘*long-tail problem*’ as described in the previous pioneering publication [20].

SH-based strategy has been widely used for protein function annotations [25-27], and many tools have been developed (*BLAST*, *FASTA*, *GoFDR*, *etc.*) [28-30], but they are severely dependent on the homology among the analyzed sequences [31-34]. To deal with this issue, ML-based strategy has thus been constructed, which learns protein function irrespective of sequence homology [35-40]. Some typical tools under this strategy include *DeepGOPlus*, *PFmulDL* & *NetGO2* [16-18], all of which apply machine learning framework(s) to achieve good annotation performance, such as the *NetGO2* in “*the 4th critical assessment of functional annotation*” (CAFA4) challenge [21]. However, due to the overwhelming domination of proteins in the *Head Label Levels* (the average number of proteins in each family of *Head Label Levels* equals to 4,210, which is ∼5 times larger than that (886) of *Tail Label Levels*, illustrated in **Fig. 1**), it is still extremely challenging for existing methods/tools to improve the prediction accuracies for families in *Tail Label Levels*, and the “*long-tail problem*” in protein functional annotation remains unsolved [41-43].

Herein, an innovative strategy, entitled ‘*AnnoPRO*’, for protein function annotation was therefore constructed. ***First***, a new method enabling image-like protein representations was proposed. This method is unique in capturing the intrinsic correlations among protein features, which can greatly favor the application of the *state-of-the-art* deep learning methods popular in image classification. ***Second***, a multimodal framework that integrates *multichannel convolutional neural network* and *long short-term memory neural network* was constructed to realize a deep learning-based protein function annotation. This framework was inspired by the *state-of-the-art* method [44] adopted in image classification for effectively dealing with its “*long-tail problem*”, *AnnoPRO* was therefore expected to greatly improve the annotation performance of the families in the ‘*Tail Label Levels*’. ***Finally***, multiple case studies based on benchmark were conducted, which further confirmed the superiority of our new method among the existing ones. All in all, *AnnoPRO* performed well and would become an essential complement to existing methods in protein function prediction.

## 2. Results and Discussion

### 2.1 Comparing the Overall Performance with Existing Tools

In this study, a total of 92,120 protein sequences were *first* collected from the competition of ‘*4th critical assessment of functional annotation*’ (CAFA4, released on Oct 21, 2019) [21], and these data were adopted to construct the annotation model (*Training* and *Validation*). *Second*, a process identical to that of ‘CAFA4’ for constructing the “*Independent Testing Dataset*” was also applied, which collected 5,623 proteins with new experimental annotation in *SwissProt* during the period from Oct 22, 2019 to May 31, 2022 [4]. As reported, the above methodology for dividing CAFA data into two datasets (one for model construction & another for independent testing) was widely adopted by previous publications [15-18] not only for developing the function annotation models, but also for making the systematic comparison among existing methods/tools.

To assess the overall performance of our new strategy, a comparison among the performances of *AnnoPRO* and eight popular methods (such as: *Diamond_BLAST_* [29], *DeepGO* [15], *DeepGOCNN* [16], *DeepGOPlus* [16], *TALE* [45], *PFmulDL* [17], *NetGO2* [18] & *NetGO3* [40]) was therefore conducted. The comparison was realized using the dataset generation approach that was provided in the above paragraph, and all methods/tools were trained on experimental function annotations that appeared before Oct 22, 2019 and tested on those annotations asserted between Oct 22, 2019 and May 31, 2022. As shown in **Table 1**, among those eight existing methods/tools, *DeepGOPlus*, *PFmulDL*, and *NetGO3* demonstrated the best performances on the GO data of BP, CC, and MF, respectively (highlighted by the *<u>underline</u>*. *Diamond_BLAST_* provided a better F*_max_* than *NetGO3* on MF data, but its AUPRC was much lower than that of *NetGO3*, therefore *NetGO3* was considered to give the best performance on MF data). These results indicated that there was no existing tool performing consistently the best under all GO classes (BP, CC, and MF). However, as shown in **Table 1**, comparing with other methods, *AnnoPRO* provided the consistently better performances (highlighted in BOLD) under all GO classes (BP, CC & MF). Particularly, when comparing with those three best performing methods/tools (*DeepGOPlus*, *PFmulDL* & *NetGO3*), the percentages of performance enhancement varied from 2.7% to 15.7% (as assessed by F*_max_*) and from 2.3% to 22.2% (as assessed by AUPRC), which illustrated a dramatical elevation in the performances of protein functional prediction by the new *AnnoPRO* strategy proposed in this study.

**Table 1.**
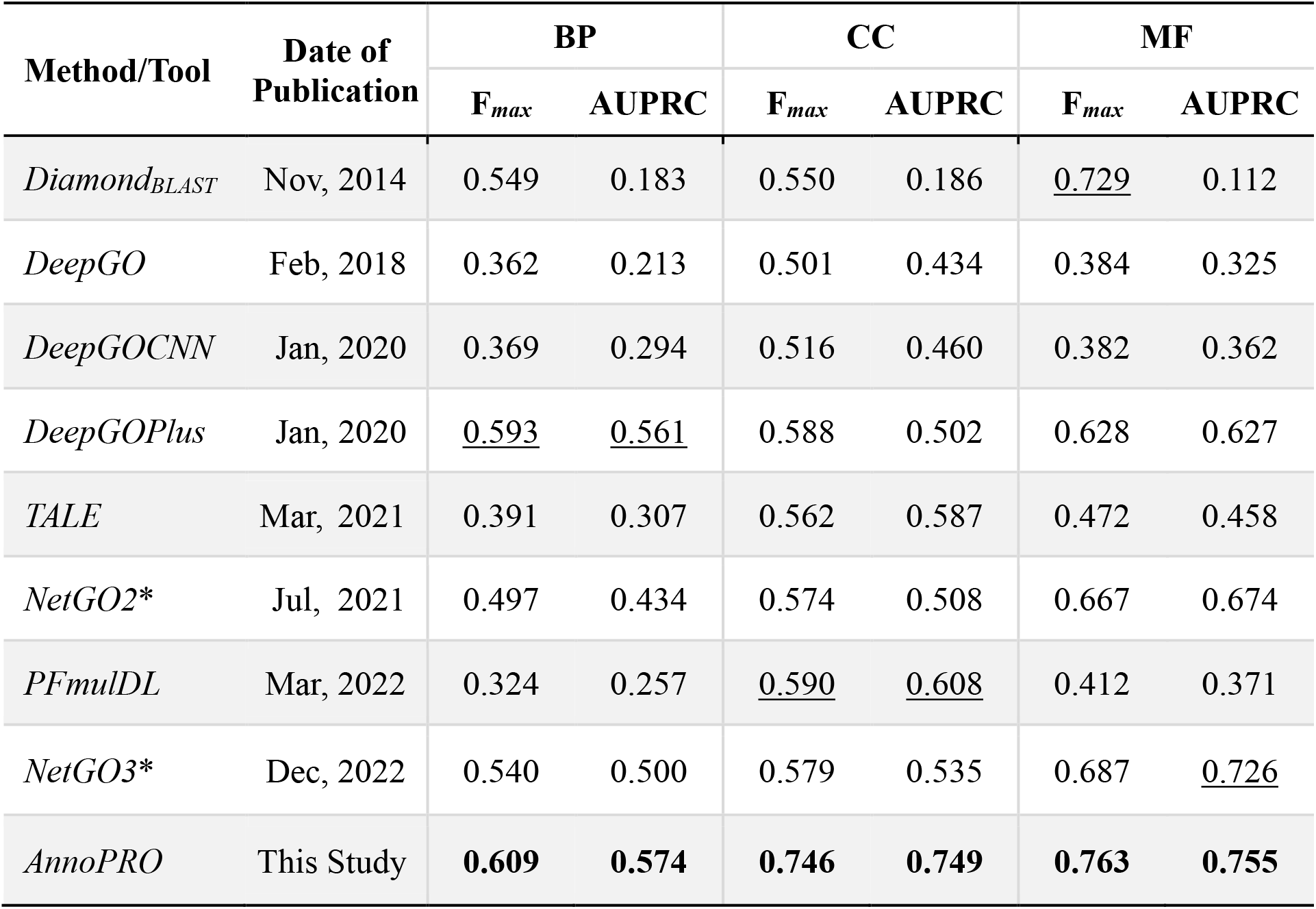
A comparison among the performances of *AnnoPRO* and eight available methods/tools. The values indicating the best performances among all methods/tools were highlighted in BOLD, and *AnnoPRO* performed consistently the best in all Gene Ontology (GO) classes (BP, CC, MF) under both evaluating criteria (F*_max_*, AUPRC). All methods/tools were ordered according to their publication dates. BP: *biological process*; CC: *cellular component*; MF: *molecular function*; F*_max_*: *protein centric maximum F-measure*; AUPRC: *area under the precision-recall curve*. Those tools marked by an asterisk (*) indicated that their source-codes for model construction were not fully provided, which made it impossible for us to train models on experimental functional annotations that appeared before Oct 22, 2019, and their performances (evaluated by F*_max_* and AUPRC) were assessed by directly uploading those experimental function annotations asserted between Oct 22, 2019 and May 31, 2022 to the online server of those annotation tools. Among those eight existing methods/tools, the best performing ones under each category were highlighted by *<u>underline</u>*.

To have an in-depth understanding on the significant elevation in the annotation performance of *AnnoPRO*, an *ablation* experiment [46] was further conducted to assess the performance changes induced by depriving some key *AnnoPRO* modules. As described in **Supplementary Fig. S1**, “<u>No *ProSIM*</u>” indicated that the *Deep Neural Network* of five fully-connected layers (5FC-DNN) was made absent from the ***Module 2*** of **Fig. 2**; “<u>No *ProMAP*</u>” indicated that the seven-channel *Convolutional Neural Network* (7C-CNN) was made absent from the ***Module 2*** of **Fig. 2**; “<u>No</u> <u>LSTM</u>” indicated that the *Long Short-Term Memory recurrent neural network* (LSTM) was made absent from the ***Module 3*** of **Fig. 2**; “<u>SC map</u>” indicated that the “*Transformation*” step in the ***Module 2*** of **Fig. 2** was deprived, and only Single-Channel (not Multi-Channel) *ProMAP* was considered; “<u>Shuffled map</u>” indicated that the *Template Map* for feature reset described in **Fig. 3*b*** was shuffled (this made the intrinsic correlation among protein features, that was captured by *ProMAP* in **Fig. 3**, completely lost). It is clear to see that the deprivation of any key *AnnoPRO* module will result in significant decrease in the annotation performance, which indicated that all the key modules collectively contributed to the good performance of *AnnoPRO*.

**Fig. 2.**
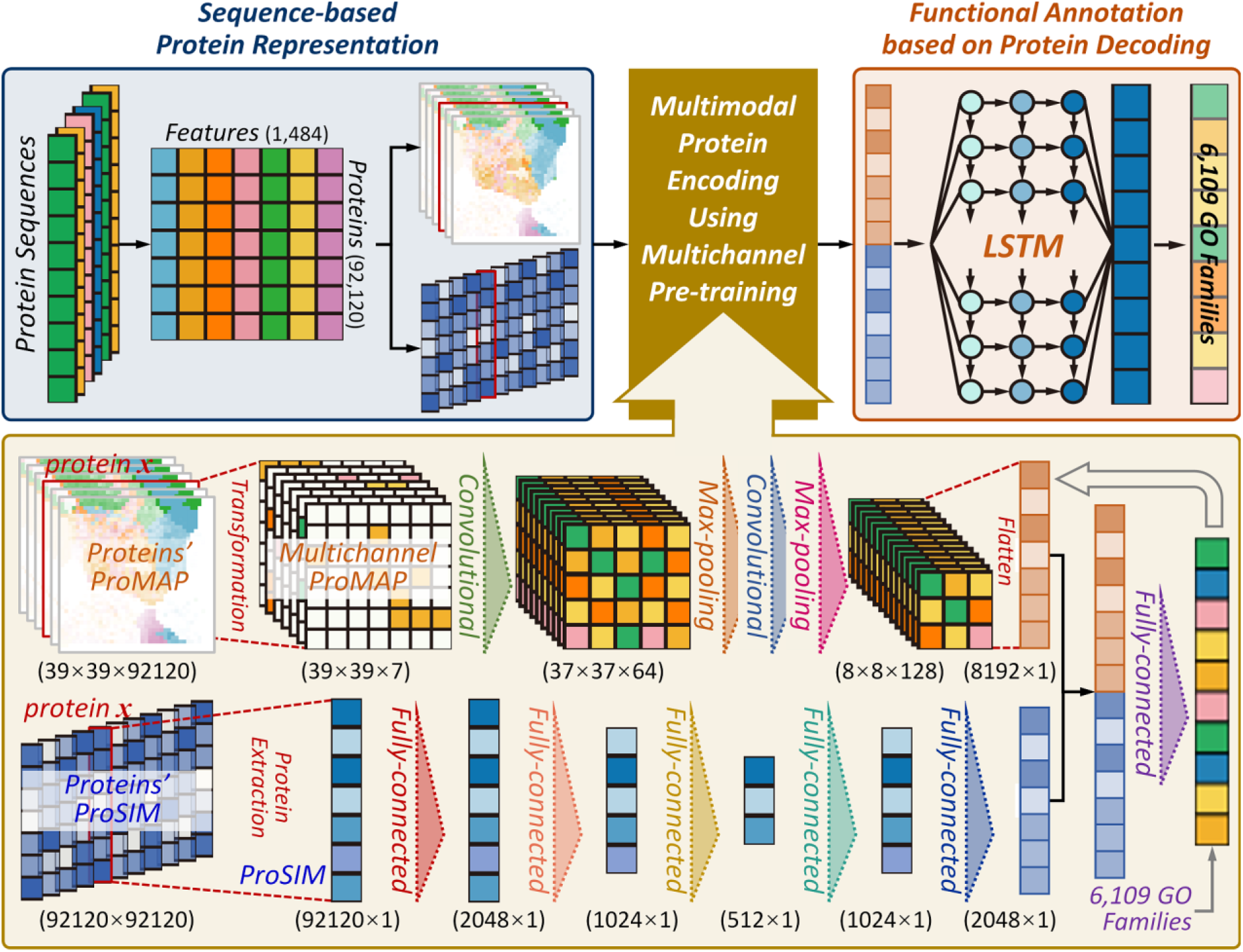
The multimodal and multichannel deep learning framework of *AnnoPRO*. There were three consecutive modules (***M1*** to ***M3***). (***M1***) sequence-based protein representation that realized the conversion of all protein sequences to the image-like protein representation. (***M2***) multimodal protein encoding using multichannel pre-training. Using the *ProMAP* and *ProSIM* generated here for proteins, a multimodal and multichannel deep learning architecture was constructed based on a seven-channel *convolutional neural network* (7C-CNN) & a *deep neural network* of five fully-connected layers (5FC-DNN) to pre-train the features of all CAFA4 proteins by integrating their annotation data of GO functional families. (***M3***) functional annotation based on protein decoding. The protein features pre-trained using the dual-path multimodal encoding layer in second module were concatenated and then fed into a *long short-term memory recurrent neural network* (LSTM) to enable a comprehensive multilabel annotation of proteins to 6,109 functional GO families.

**Fig. 3.**
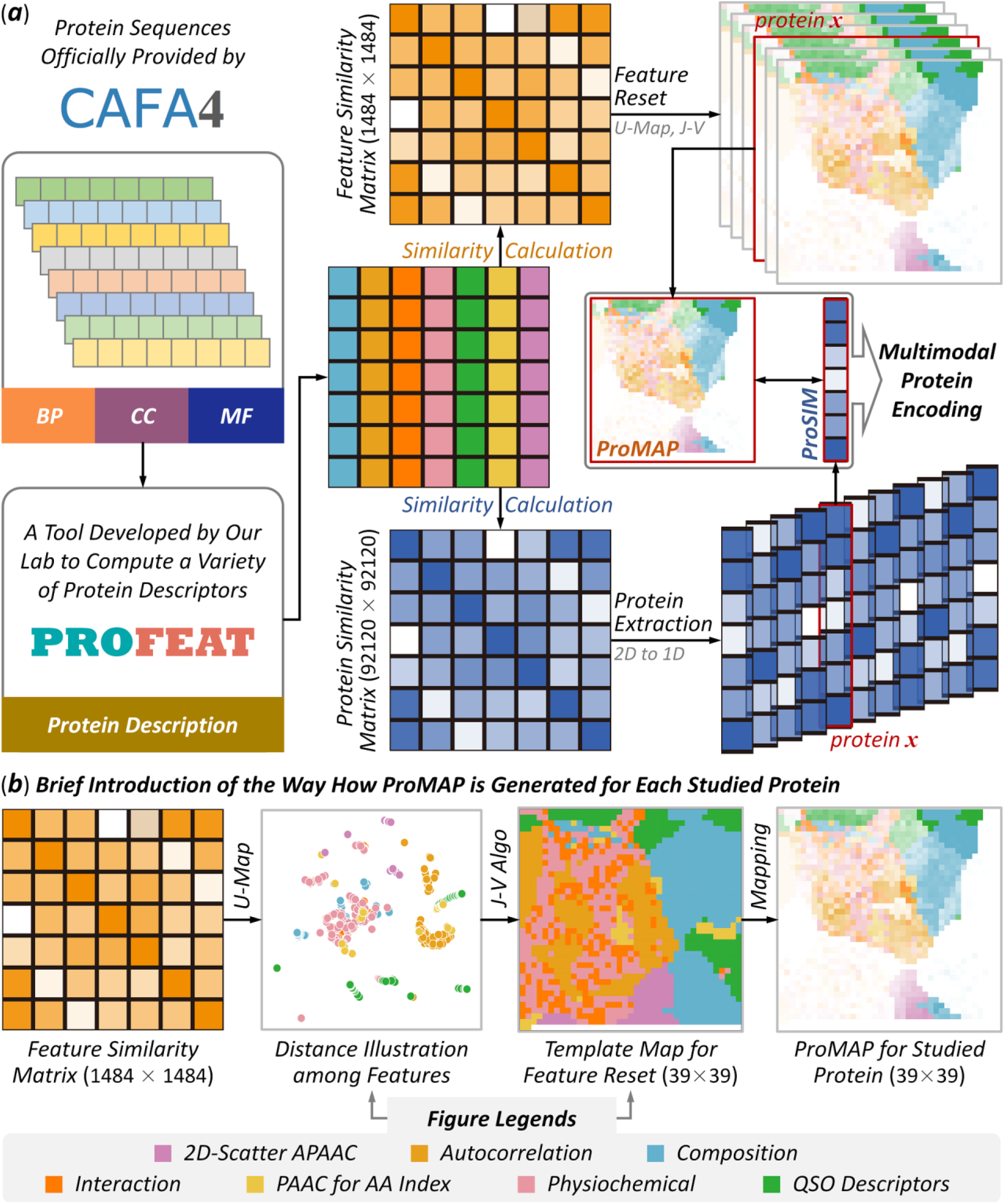
A schematic illustration of the procedure used in this study facilitating sequence-based protein representation. (***a***) a concrete description on the novel strategy proposed in this study for representing protein sequences based on calculating the similarities among proteins and features. Particularly, a total of 92,120 protein sequences were *first* collected from the official website of CAFA4, and their descriptors were computed by a popular tool developed by our research lab [60]; two similarity matrices (showing similarity among features & proteins) were *then* generated; the feature similarity matrix was *further* applied to reset the location of all features in a 2D map, and any sequence could therefore be represented as a 39×39 square image (which was named in this study as ‘*ProMAP*’ for each protein); the protein similarity matrix was *also* adopted to extract a 1D vector, and any protein sequence could be represented as a vector of 92,120 features (which was named here as ‘*ProSIM*’ for each protein); and both *ProMAP* and *ProSIM* were *finally* used to facilitate the subsequent “*multimodal protein encoding*” (the second module shown in **Figure 2**). (***b***) a brief introduction of the way how a ‘*ProMAP*’ was created for each protein. Particularly, feature similarity matrix was *first* applied to illustrate the distances among any two features based on U-Map [61]; a template image/map for feature reset was *then* generated by *Jonker-Volgenant* (*J-V*) algorithm [62] and U-Map; and *ProMAP* was *finally* produced for each protein by mapping the intensities of all features to their corresponding locations in the template image.

### 2.2 Level-based Performance Comparison with Existing Tools

Based on those analyses above, three recently-published methods/tools (*DeepGOPlus*, *PFmulDL* & *NetGO3*) were identified to perform better than the others, and were reported as “*state-of-the-art*” by previous publication [47]. Therefore, a comparison among the level-based performances of *AnnoPRO* and these three *state-of-the-art* methods was conducted. The so-called level-based performances were based on the hierarchical structure of GO families provided in the first section of **Materials and Methods** and the definition in **Supplementary Fig. S2**. As shown in **Fig. 4**, the level-based performances were represented using the AUC value in predicting testing data, and the performances of *AnnoPRO*, *DeepGOPlus*, *NetGO3* & *PFmulDL* were highlighted using light red, light green, orange & light blue, respectively (all values were shown in **Supplementary Table S1**). For the GO families in ‘*Head Label Level*’ (LEVEL 2 & LEVEL 3 in **Supplementary Fig. S2**), the performance of *AnnoPRO* was roughly as good as that of the other three methods (1.4∼4.1% improvements in most cases, but 0.1% decline in one single case). For the GO families in ‘*Tail Label Level*’ (LEVEL 4 to LEVEL 10 in **Supplementary Fig. S2**), *AnnoPRO* showed the consistently superior performance among all methods (1.7∼28.2% improvements in all cases). Particularly, 13 (61.9%) out of all 21 improvements were higher than 5%, and 6 (28.6%) out of those 21 improvements were larger than 10% (as illustrated in **Fig. 4**).

**Fig. 4.**
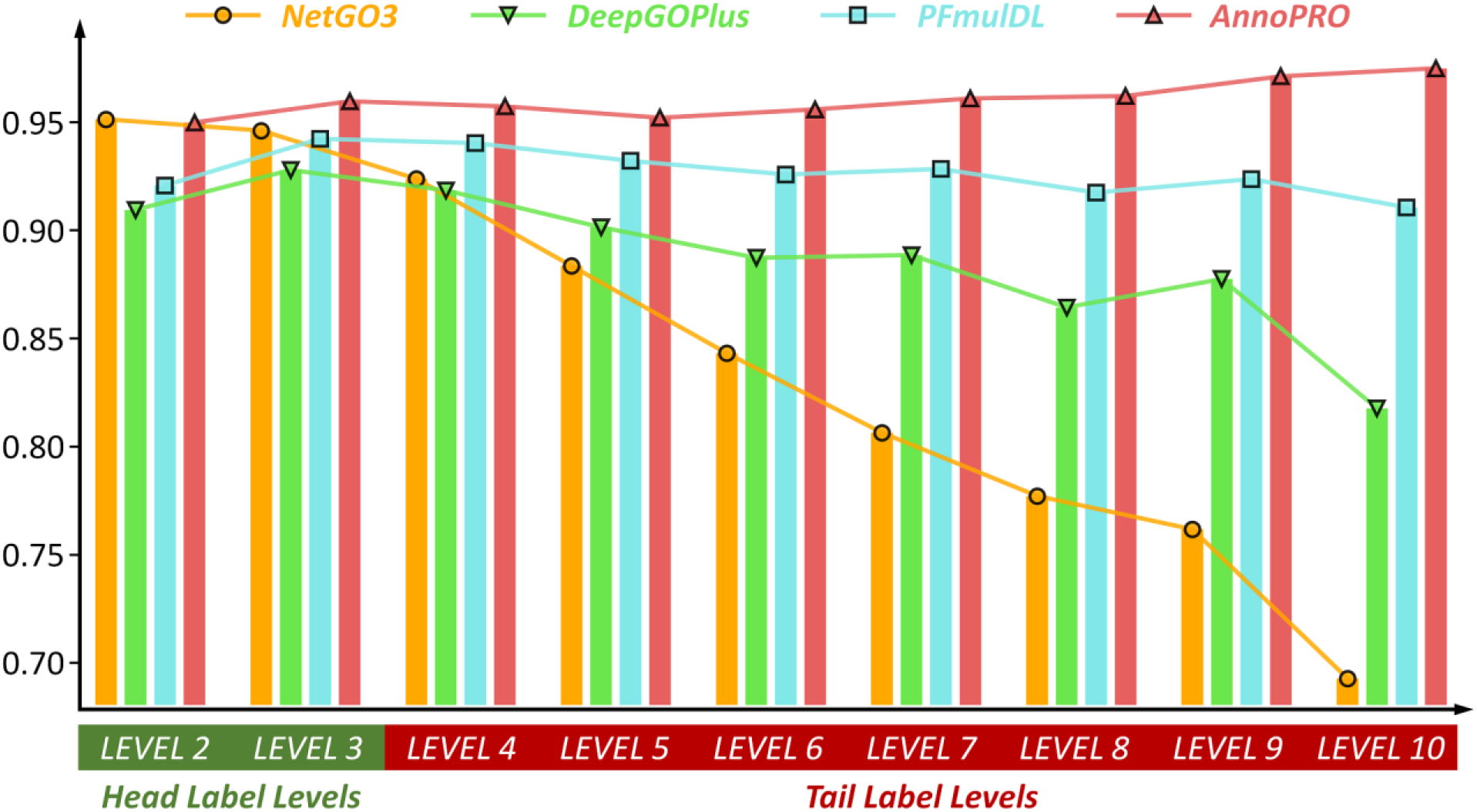
A comparison among the performances of *AnnoPRO* and three representative methods. The performances were represented using AUC values in predicting the experimentally validated new protein functions that were not included in CAFA4 data, and the performances of *AnnoPRO*, *DeepGOPlus*, *NetGO3* and *PFmulDL* were highlighted in light red, light green, orange and light blue, respectively. For the GO families in the ‘*Head Label Levels*’ (LEVEL 2 & LEVEL 3 shown in **Supplementary Fig. S2**), the performance of *AnnoPRO* was roughly as good as that of the other three methods (1.4∼4.1% improvements in most cases, but 0.1% decline in one single case). For the GO families in the ‘*Tail Label Levels*’ (LEVEL 4 to LEVEL 10 shown in **Supplementary Fig. S2**), *AnnoPRO* demonstrated the consistently superior performance among four methods (1.7∼28.2% improvements in all cases). Particularly, 13 (61.9%) out of all 21 improvements were over 5%, and 6 (28.6%) out of 21 improvements were more than 10%. Therefore, *AnnoPRO* was identified *superior* in significantly improving the annotation performances of the families in ‘*Tail Label Levels*’ without sacrificing that of the ‘*Head Label Levels*’, which was highly expected to make contribution to solving the long-standing ‘*long-tail problem*’ [20] in functional annotation.

Furthermore, as illustrated in **Fig. 4**, *DeepGOPlus* and *NetGO3* performed well in LEVEL 2 and LEVEL 3, but experienced a dramatic decline of performance from LEVEL 4 to LEVEL 10. This clearly showed that the “*long tail problem*” remained a serious issue for the protein function annotation using existing methods (significantly declined from 95.1% to 69.3% for *NetGO3* and from 91.8% to 81.8% for *DeepGOPlus*). The *PFmulDL* was a method that could largely enhance the performances for the GO families in ‘*Tail Label Level*’, but *AnnoPRO* provided a much better performances in all levels than *PFmulDL* (as shown in **Fig. 4**). In other words, *AnnoPRO* was the first method reported to achieve *superior* performance in protein annotations for GO families in ‘*Tail Label*’ levels without sacrificing that in ‘*Head Label*’ ones, which was therefore expected to highly contribute to the final solution of the long-standing ‘*long-tail problem*’ [20].

### 2.3 Performance Comparison Using Proteins from Various Species

Sequence variation among the orthologs of various species may induce subtle, or even substantial, changes in protein structure, which may lead to proteins with similar sequence showing different functions [48-50]. This leads to great difficulty in functional annotations for orthologous proteins [51], and it is therefore of great interests to compare the capacities of *AnnoPRO* and the *state-of-the-art* methods/tools (*DeepGOPlus*, *PFmulDL* and *NetGO3*) from this perspective. In this study, the species origins of 92,120 proteins from CAFA4 (adopted as ‘*Training*’ & ‘*Validation*’ datasets for developing the annotation model in this study) were ***<u>first</u>*** analyzed, and 17 species were found (*homo sapiens*, *mus musculus*, *drosophila melanogaster*, *arabidopsis thaliana*, *etc.*). Meanwhile, the species origins of 5,623 proteins (adopted as ‘*Independent Testing*’ dataset in this study) were also found, which discovered a total of 1,014 species (despite those 17 species, there were many other species: *bos taurus*, *camellia sinensis*, *canis lupus familiaris*, *gallus gallus*, *mycobacterium tuberculosis*, *oryza sativa*, *etc.*). ***<u>Second</u>***, the 5,623 proteins were further divided into two groups. One group included 1,859 proteins (titled ‘**SameSP**’) from those 17 species covered by ‘*Training*’ & ‘*Validation*’ datasets, and another had 3,764 proteins (titled ‘**DiffSP**’) from the remaining 997 species unique in ‘*Independent Testing*’ dataset. ***<u>Third</u>***, the performances of *AnnoPRO* and those two *state-of-the-art* methods/tools (*DeepGOPlus* and *PFmulDL*; *NetGO3* was not included here since its source code for model construction was not accessible) were evaluated based on the two groups of ‘*Independent Testing*’ data, and the evaluating results were provided in **Table 2**.

**Table 2.**
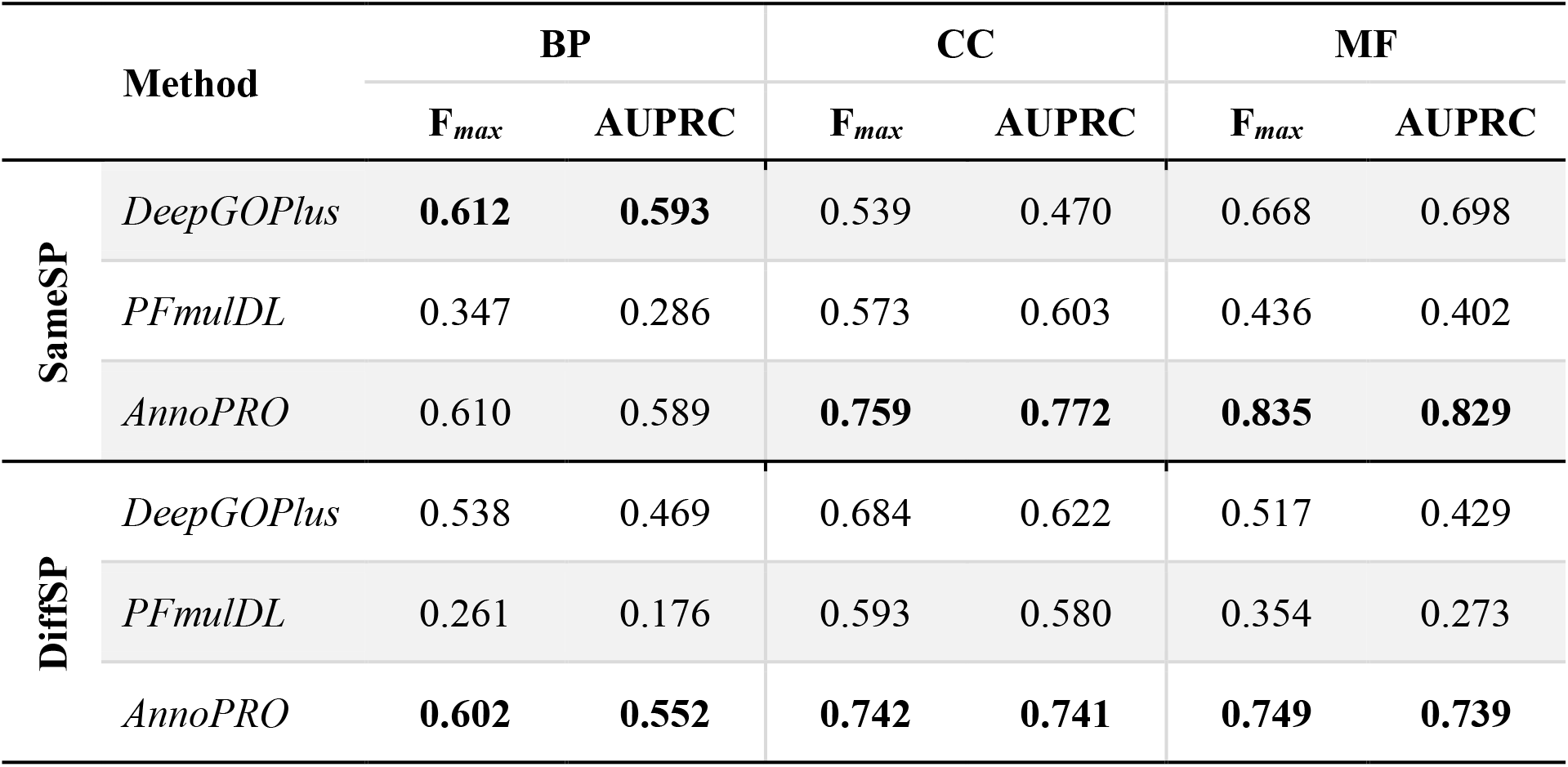
A comparison among the performances of *AnnoPRO* and two *state-of-the-art* methods (*DeepGOPlus* and *PFmulDL*) on predicting two groups of ‘*Independent Testing*’ data (**SameSP** and **DiffSP**). **SameSP** had 1,859 proteins from 17 species covered by ‘*Training*’ and ‘*Validation*’ datasets of this study; **DiffSP** included 3,764 proteins from the remaining 997 species unique in ‘*Independent Testing*’ data of this study. Those values indicating the best performance among all three methods were highlighted in BOLD, and *AnnoPRO* performed the best in the vast majority of the Gene Ontology (GO) classes (BP, CC & MF) under both evaluating criteria (F*_max_*, AUPRC). BP: *biological process*; CC: *cellular component*; MF: *molecular function*.

As shown in **Table 2**, the *AnnoPRO* performed the best in the vast majority of the Gene Ontology classes (BP, CC, MF) under both evaluating criteria (F*_max_*, AUPRC), and those values indicating the best performance among those three methods (*AnnoPRO*, *DeepGOPlus*, and *PFmulDL*) were highlighted in BOLD. Particularly, for the **SameSP** group of *Independent Testing* data, *AnnoPRO* showed superior performance in both CC and MF with significant elevations in F*_max_* and AUPRC (elevated by 0.13 to 0.43), and *AnnoPRO* demonstrated equivalent performance in BP comparing with *DeepGOPlus* with slightly lower F*_max_* and AUPRC (lower by 0.002 and 0.004, respectively); for the **DiffSP** group of data, the performances of *AnnoPRO* remained the best in CC & MF with significant elevation in F*_max_* and AUPRC (elevated by 0.06 to 0.47), and the *AnnoPRO* performed better in the BP comparing with *DeepGOPlus* (F*_max_* and AUPRC were elevated by 0.06 and 0.08). All in all, the results indicated that *AnnoPRO* gave good predictive performances on *independent* data whose species origins were covered by *training*-*validation*, and its predictive performances on *independent* data whose species origins were distinct from that of *training*-*validation*, became even better when comparing with *state-of-the-art* methods. In other words, the *AnnoPRO* showed good capacity on predicting the proteins that have little representativeness in *training*-*validation* data, which was very valuable for the function annotation of novel proteins from the species not covered by both ‘*Training*’ and ‘*Validation*’ datasets during model construction.

### 2.4 Functional Annotation of the Homologous Proteins with Distinct Functions

As reported, a small variation in sequence could lead to vastly different functional outcomes [52], which made the annotation of homologous proteins with distinct functions a great challenge and a fascinating direction for the researchers in related research community. In order to evaluate the predictive performances of *AnnoPRO* and three *state-of-the-art* methods on such kind of proteins, two pairs of homologous proteins of distinct functions were then analyzed: *growth differentiation factors* (GDF8 & GDF11) and *heat shock proteins* (HSPA1A & HSPA2).

#### 2.4.1 Case Study 1 on Different Growth Differentiation Factors

*Growth differentiation factors* (GDFs) belong to the transforming growth factor β (TGFβ) family, which regulate the formation of central nervous system [53]. GDF11 is a protein in the family of GDFs, which shares 65% sequence similarity with GDF8 (*myostatin*, MSTN) and 90% sequence identity in the active domain [54]. As reported, the interaction between GDF8 and *follistatin-288* (FS288) formed a complex to bind heparin, which defined the molecular mechanisms underlying GDF8’s key GO family: ‘*heparin binding*’ (GO:0008201) [55]. Different from GDF8, the varied residues in GDF11 made it unable to interact with FS288, and it therefore suffered from the loss of ‘*heparin binding*’ function [56]. The sequences between GDF8’s and GDF11’s active domains were aligned in **Fig. 5*a***, where varied residues between two GDFs were marked in light green and blue background, respectively. Combined with the structural superimpositions (illustrated in **Fig. 5*b***) between GDF8 (light green) & GDF11 (blue) and their interactions with FS288 (gray surface) [57], three varied residue pairs (F315Y, V316M & L318M located in the binding surface between GDF and FS288) were found as key residues for ‘*heparin binding*’ function [58].

**Fig. 5.**
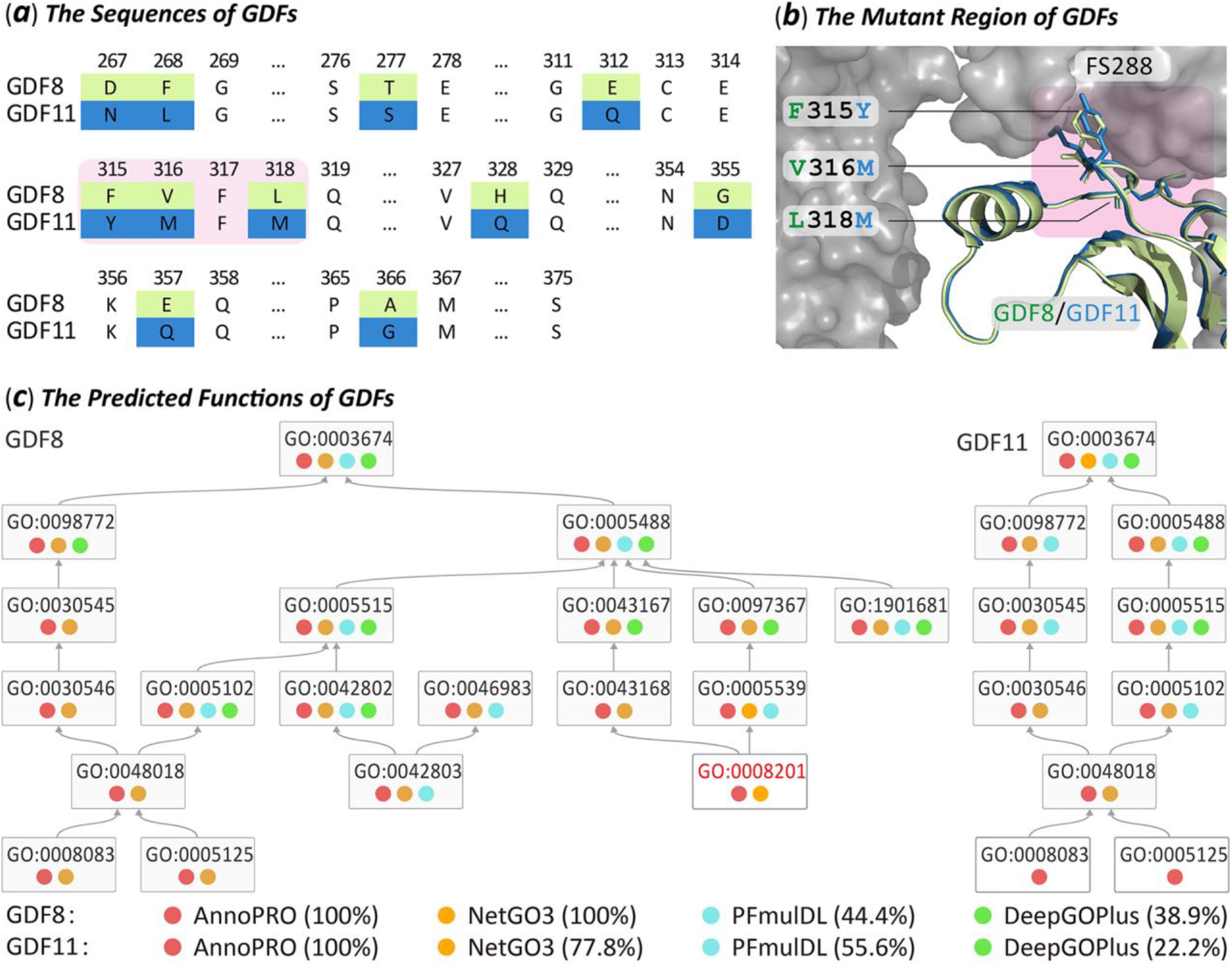
Performance assessment of four methods using two well-known *growth differentiation factors* (GDF8, GDF11). As reported, the interaction between GDF8 and follistatin-288 (FS288) formed a protein complex to bind ‘heparin’, which defined the molecular mechanisms underlying GDF8’s key GO family: ‘*heparin binding*’ (GO:0008201) [55]. Different from GDF8, the varied residues in GDF11 made it unable to interact with FS288, and it therefore suffered from the loss of the ‘*heparin binding*’ function [56]. (***a***) sequence alignment between GDF8 and GDF11, where varied residues between two GDFs were marked in light green and blue background, respectively. Three varied residue pairs (F315Y, V316M & L318M on the binding surface between GDF8 and FS288) which were found as key residue indicating GDFs’ ‘*heparin binding*’ function [58], were shown in pink background. (***b***) structural superimposition between GDF8 (light green) & GDF11 (blue) and their interactions with FS288 (gray surface). As highlighted in pink background, those three key residue pairs (F315Y, V316M & L318M) located right in the binding interface between GDF and FS288. (***c***) the results of functional annotation predicted by four studied methods. If a GO family is successfully predicted by a method, a colored circle will be adopted to indicate the prediction result. Particularly, a successful prediction made by *AnnoPRO*, *NetGO3*, *PFmulDL* or *DeepGOPlus* was indicated by a circle of light red, orange, light blue or light green, respectively. As shown, *AnnoPRO* is the only one that can successfully predict all GO families for both GDFs.

In this study, the ‘*heparin binding*’ function (GO:0008201) for the wild type GDF8 (***GDF8-WT***) and its two mutants (***GDF8-Mutant-1*** and ***GDF8-Mutant-2***) was predicted using *AnnoPRO* and three *state-of-the-art* tools (*DeepGOPlus*, *PFmulDL*, *NetGO3*). ***GDF8-Mutant-1*** contains eight mutations (D267N, F268L, T277S, E312Q, H328Q, G355D, E357Q & A366G) which locate far away from the binding interface between GDF8 and FS288. The interaction between ***GDF8-WT*** and FS288 forms a complex binding with heparin, which is the molecular mechanism underlying ***GDF8-WT***’s ‘*heparin binding*’ function (GO:0008201). Since all eight mutations were far away from the binding interface between GDF8 & FS288, it is expected that ‘*heparin binding*’ function remains in ***GDF8-Mutant-1*** [58]. Meanwhile, ***GDF8-Mutant-2*** contains three mutations (F315Y, V316M & L318M, on the bindin *binding* function [58]. In other words, it is expected that ***GDF8-Mutant-2*** loses its wild type’s ‘*heparin binding*’ function [58]. All in all, there is gain-of-function of ‘*heparin binding*’ in ***GDF8-WT*** and ***GDF8-Mutant-1***, while there is loss-of-function in ***GDF8-Mutant-2***. As described in **Table 3**, ‘Success’ denoted that the gain/loss-of-function is successfully predicted by method, while ‘Fail’ showed that the prediction by method is incorrect. As shown, *AnnoPRO* was the only method that “successfully” captured the significant functional variations induced by small amount of residue mutations among GDF8 proteins.

**Table 3.**
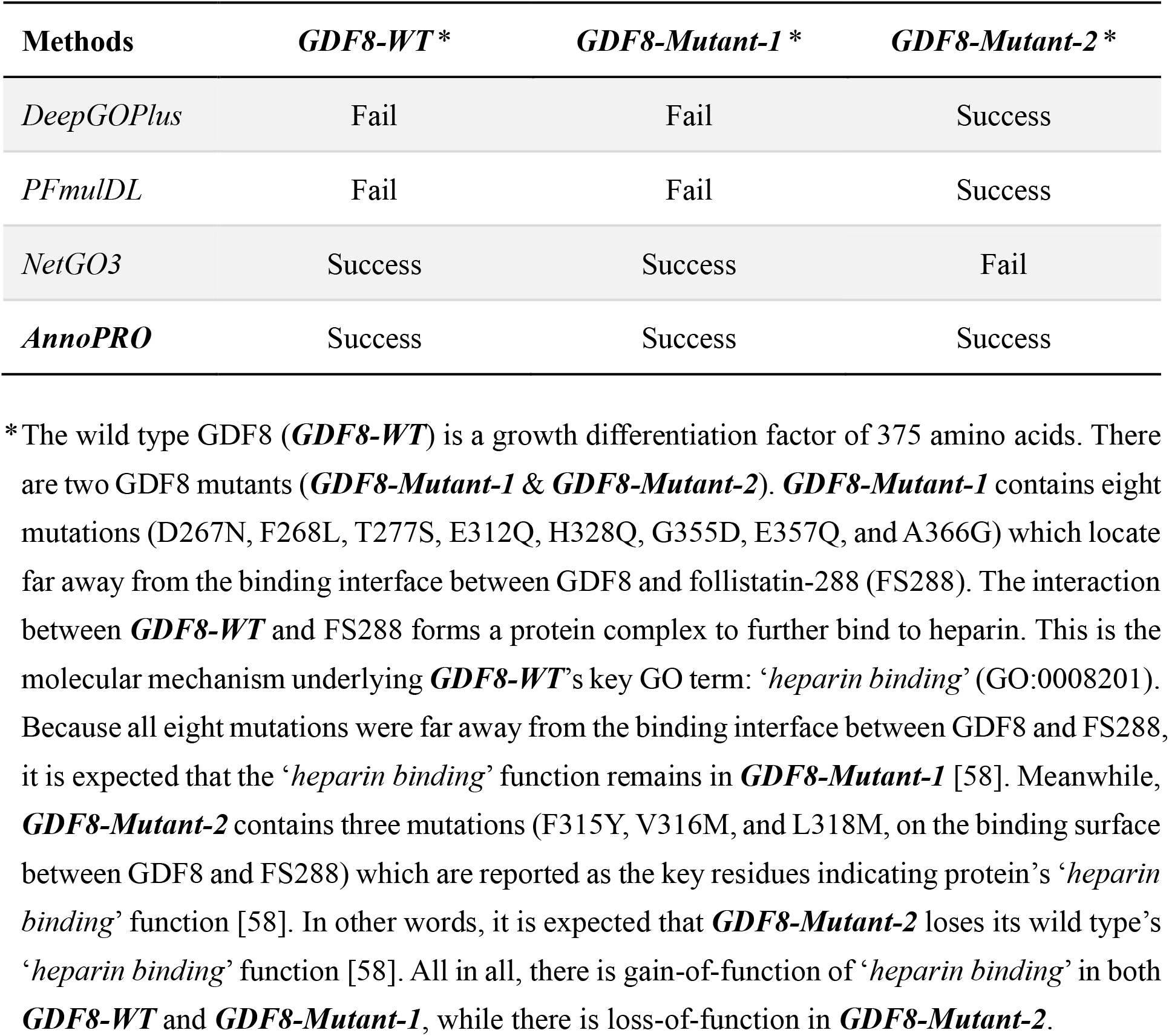
The prediction of the ‘*heparin binding*’ function (GO:0008201) for the wild type GDF8 (***GDF8-WT***) and its two GDF8 mutants (***GDF8-Mutant-1*** & ***GDF8-Mutant-2***) using *AnnoPRO* and three representative methods. ‘Success’ denotes that the gain/loss-of-function is successfully predicted by the corresponding method, while ‘Fail’ indicates that it is incorrectly predicted. As demonstrated, significant functional variations among ***GDF8-WT***, ***GDF8-Mutant-1***, and ***GDF8-Mutant-2*** can only be “successfully” captured by our newly developed AnnoPRO.

Moreover, the sequences of GDF8 and GDF11 were reported to be highly homologous, but their functions were distinct with 291 different GO families. Therefore, it was of great interests to test the predictive performances of *AnnoPRO* and three *state-of-the-art* tools on this issue. As shown in **Table 4**, *AnnoPRO* performed the best in the vast-majority (11/12) of the Gene Ontology (GO) classes (BP, CC, MF) under both evaluating criteria (recall & precision). Taking the GO class of MF as an example (as shown in **Fig. 5*c***), GDF8 and GDF11 contained 19 and 10 MF families, respectively, and the functions annotated by those four methods were highlighted. If a MF family is successfully predicted by method, a colored circle will be used to indicate the prediction result. As illustrated in **Fig. 5*c***, the successful prediction made by *AnnoPRO*, *NetGO3*, *PFmulDL* or *DeepGOPlus* was indicated by a circle of light red, orange, light blue or light green, respectively, and *AnnoPRO* is the only one that can successfully predict all MF families for both GDFs.

**Table 4.**
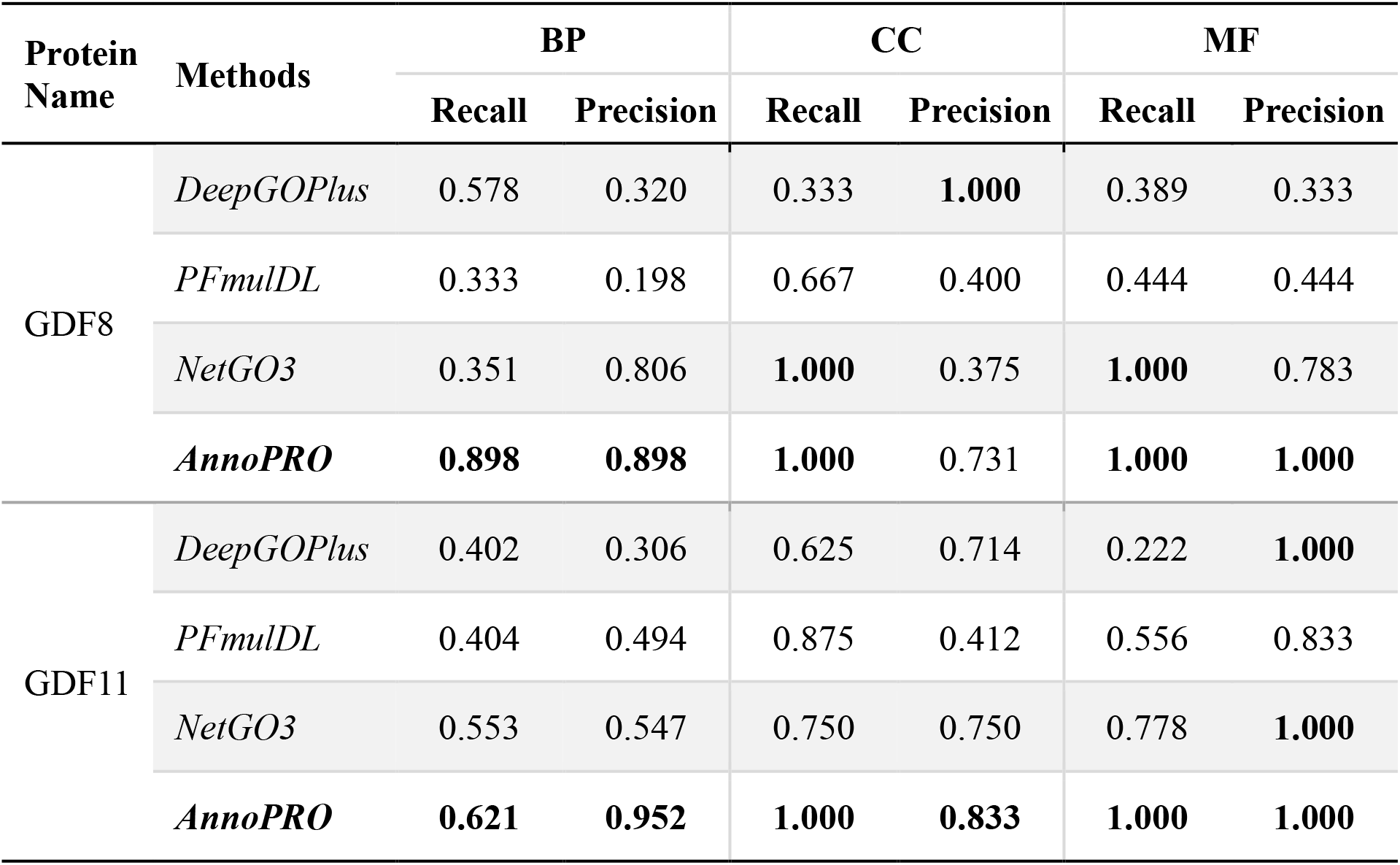
A comparison among the predictive performances of *AnnoPRO* and three representative methods for the functional annotations of two well-known *growth differentiation factors* (GDF8, GDF11). Those values indicating the best performances among all methods were highlighted in BOLD, and *AnnoPRO* performed the best in the vast-majority (11/12) of the Gene Ontology (GO) classes (BP, CC, MF) under both evaluating criteria (recall, precision). All methods were ordered based on their publication dates. BP: *biological process*; CC: *cellular component*; MF: *molecular function*; GDF8: *growth differentiation factor 8*; GDF11: *growth differentiation factor 11*.

**Table 5.**
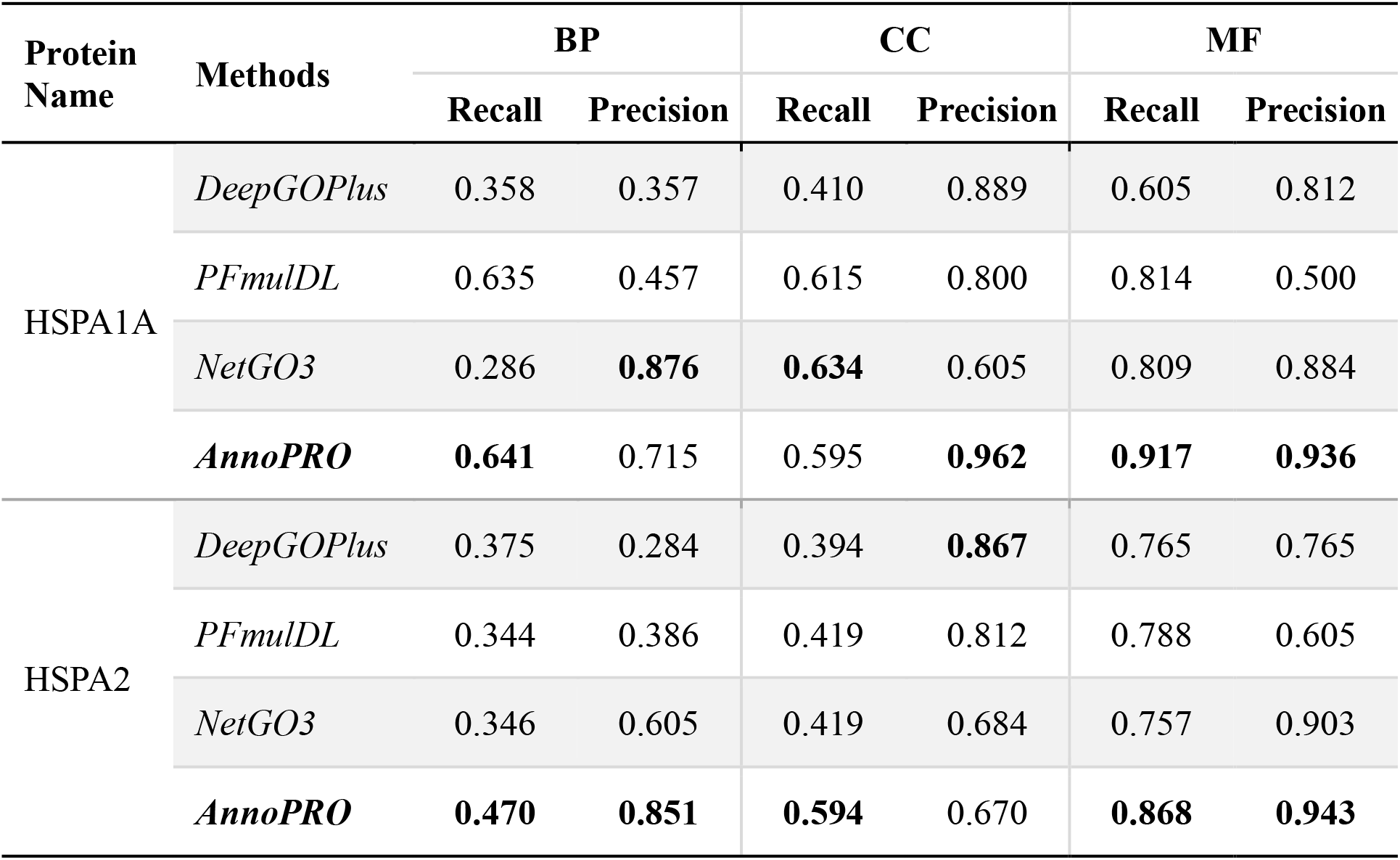
A comparison among the predictive performances of *AnnoPRO* and three representative methods for the functional annotations of two well-known *heat shock 70kDa proteins* (HSPA1A, HSPA2). Those values indicating the best performances among all methods were highlighted in BOLD, and *AnnoPRO* performed the best in the vast-majority (9/12) of the Gene Ontology (GO) classes (BP, CC, MF) under both evaluating criteria (recall, precision). All methods were ordered based on their publication dates. BP: *biological process*; CC: *cellular component*; MF: *molecular function*; HSPA1A: *heat shock 70 kDa protein 1A*; HSPA2: *heat shock 70 kDa protein 2*.

#### 2.4.2 Case Study 2 on Different Heat Shock Proteins

*Heat shock proteins* (HSPs) are ubiquitous and conserved proteins in prokaryotic and eukaryotic organisms, which are essential for maintaining cellular proteostasis [59]. In this study, two well-known *heat shock 70kDa proteins* (HSPA1A & HSPA2) were analyzed. The sequence of HSPA2 is >80% similar to HSPA1A, while the total number of different GO families between these two proteins is >300. Thus, it was of great interest to assess the predictive performances of *AnnoPRO* and three *state-of-the-art* tools on this issue. As shown in **Table 4**, *AnnoPRO* performed the best in the vast-majority (9/12) of the GO classes under both evaluating criteria (recall and precision). Taking the GO class of MF as an example (illustrated in **Supplementary Fig. S3** for HSPA1A and **Supplementary Fig. S4** for HSPA2), the HSPA1A and HSPA2 had 44 and 35 MF families, respectively, and the functions annotated by those four methods were highlighted. If a MF family is successfully predicted by method, a colored circle will be used to indicate the prediction result. As illustrated, the successful prediction made by *AnnoPRO*, *NetGO3*, *PFmulDL* or *DeepGOPlus* was indicated by a circle of light red, orange, light blue or light green, respectively, and *AnnoPRO* is the only one that reach >90% accuracies in predicting MF families for both HSPs. Furthermore, there were 16 different MF families between HSPA1A and HSPA2 (highlighted by red frames in **Supplementary Fig. S3** for HSPA1A and **Supplementary Fig. S4** for HSPA2). As shown, *AnnoPRO* performed the best in most (13/16) families, and *NetGO3*, *PFmulDL* and *DeepGOPlus* successfully predicted 7, 10 and 3 families, respectively.

## 3. Materials and Methods

### 3.1. The Collection of Benchmark Dataset for the Construction of *AnnoPRO*

In this study, a total of 92,120 protein sequences were *first* collected from the competition of ‘*4th critical assessment of functional annotation*’ (CAFA4, released on Oct 21, 2019) [21], and these data were adopted to construct the annotation model (*Training* and *Validation*). *Second*, a process identical to that of ‘CAFA4’ for constructing the “*Independent Testing Dataset*” was also applied, which collected 5,623 new protein annotations manually reviewed in *SwissProt* during the period from Oct 22, 2019 to May 31, 2022 [4]. *Third*, the biological functions (denoted by GO families) of all these proteins collected above were matched directly from the UniProt knowledgebase [4]. Like existing tools [15-18], only those GO families with relatively large number of proteins (>50) were included into the model construction process of this study, which contained a total of 6,109 non-repetitive GO families. The full relations among these families were also downloaded [19].

Within the downloaded files, GO families were provided in a hierarchical structure. As illustrated in **Supplementary Fig. S2**, three root families were provided at the top of the structure, which included *biological process* (BP), *molecular function* (MF), and *cellular component* (CC). Then, the remaining GO families were hierarchically connected to the three root ones. In this study, the level of those root families was defined as ‘LEVEL 1’ (as shown in **Supplementary Fig. S2**). The direct child families of the root ones were classified to LEVEL 2, and the families of LEVEL 3 were determined by the direct child families of LEVEL 2. The following levels can be therefore deduced in the same manner. Based on our comprehensive evaluation on all GO data, the bottom level of GO’s hierarchical structure was LEVEL 11, which had no child family and composed of the smallest number of proteins comparing with the families in other levels (LEVEL 1 to 10). As shown in **Fig. 1**, the average numbers of proteins (ANP) in GO families of nine levels (LEVEL 2 to LEVEL 10) were provided. There was a clear descending trend of ANPs from LEVEL 2 to LEVEL 10. Since the ANP of one family indicated its representativeness among all families, this denoted a gradual decrease of the representativeness of a family with the penetration into deeper level. Thus, these nine levels could be classified into two groups based on their ANPs: the “*Head Label Levels*” (ANP of their GO families ≥ 2,000) and the “*Tail Label Levels*” (ANP of their GO families < 2,000). As shown, the total number (5,323) of families in “*Tail Label Levels*” was over 10 times larger than that (459) of the “*Head Label Levels*”, and such data distribution was typical for any research studies that were suffering from the ‘*long-tail problem*’ [15-18].

### 3.2. A New Multimodal Deep Learning Framework for Function Annotation

#### 3.2.1. Three Consecutive Modules Integrated in the Framework

In this study, a multimodal deep learning framework was proposed to enable function annotation, which was composed of three consecutive modules. As shown in **Fig. 2**, the three key modules included: (**1**) sequence-based protein representations that convert protein sequences to an image-like representation, (**2**) multimodal protein encoding based on multichannel pre-training, and (**3**) protein function annotations based on protein decoding. The detailed descriptions on these three consecutive modules were explicitly discussed as follows.

##### Module 1. A Novel Method Proposed to Enable Image-like Protein Representation

A novel method enabling image-like protein representation was proposed in this work, which is unique in capturing the intrinsic correlations among protein features. The adoption of this method can greatly favor the applications of the *state-of-the-art* deep learning approach popular in image classification. As shown in **Fig. 3*a***, the descriptors of all protein sequences collected from the website of CAFA4 were ***<u>first</u>*** computed using a popular method developed by our lab [60], which resulted in a total of 1,484 protein descriptors. These descriptors could be grouped into 7 classes: *amphiphilic pseudo amino acid composition*, *amino acid composition*, *physicochemical property*, *amino acid autocorrelation*, *molecular interaction*, *quasi-sequence-order* & *pseudo amino acid composition* (the detailed descriptions on each class were provided in **Supplementary Table S2**). ***<u>Second</u>***, a new protein-descriptor matrix (92,120 proteins × 1,484 features) was created, and any original number (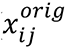) in this matrix was further normalized to 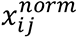 by min-max method:

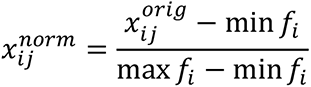

where *f*_*i*_ denoted the *ith* feature, min *f*_*i*_ and max *f*_*i*_ indicated the minimum and maximum value of the *ith* feature among all proteins, respectively. ***<u>Third</u>***, 2 similarity matrices (showing similarity among features & proteins) were generated. Taking the *feature similarity matrix* (FSM) as an example, it was produced by calculating the pair-wise distances among 1,484 features (each feature, like *f*_*a*_ & *f*_*b*_, was shown by a vector of 92,120 length) using the following equation:

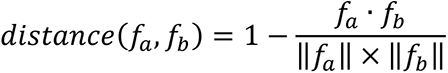

***<u>Fourth</u>***, the FSM was then adopted to reset the location of all features in a 2D map, and an image (named as ‘*ProMAP*’ in this study) was generated for each protein. This is the key step for image-like protein representation, which was explicitly described in **Fig. 3*b***. As illustrated, the FSM was *first* used to illustrate the distances among any two features based on U-Map [61]; a template image for feature reset was *then* generated by *Jonker-Volgenant* (*J-V*) algorithm [62] and U-Map; and *ProMAP* was *finally* produced for each protein by mapping the intensities of all features to their corresponding locations in the template image. As a result, the newly created *ProMAP* for each protein is unique in capturing the intrinsic correlation among diverse protein features, which facilitate the application of any deep learning methods popular in image classification. Moreover, another similarity matrix named *protein similarity matrix* (PSM) was *also* adopted to extract 1D vector for each protein, and any protein sequence could thus be represented as a vector of 92,120 features (named as ‘*ProSIM*’ in this study). Both *ProMAP* and *ProSIM* can be further applied for “*multimodal protein encoding*” (the *Module 2* shown in **Fig. 2** and discussed below).

##### Module 2. A New Multimodal Protein Encoding Using Multichannel Pre-training

In this module, a novel multimodal framework integrating a *seven-channel convolutional neural network* (7C-CNN) and a *deep neural network of five fully-connected layers* (5FC-DNN) to pre-train the features of protein was adopted. This pre-train process was expected to be very effective in transferring functional family information for optimizing the concatenated protein features [63], which could greatly facilitate the application of the *long short-term memory* (LSTM) *neural network* for function annotation in the next module [64]. Particularly, as illustrated in **Fig. 2**, the *ProMAPs* (39 × 39) for 92,120 proteins were transformed to 7 images of multichannel based on the different classes of protein descriptor, and multiple convolutional and max-pooling layers were used for learning the protein functions; the *ProSIMs* (92,120 × 1) for 92,120 proteins were extracted from the *protein similarity matrix* (PSM), and a *neural network of five fully-connected layers* (5FC-DNN) was applied for encoding proteins. By concatenating those two vectors from *ProMAP* and *ProSIM*, a total of 92,120 concatenated protein encoding vectors were created, and a fully-connected layer was further applied to refine the protein encoding by comparing with the 6,109 GO function families well-defined in *Gene Ontology*. As a result, 92,120 protein encodings were pre-trained, which could then be fed into LSTM for multilabel functional annotation.

##### Module 3. Protein Decoding-based Functional Annotation Using LSTM Method

In this module, the *long short-term memory neural network* (LSTM) was used to decode proteins for annotating their functions. LSTM had been utilized to cope with “*long-tail problem*” in *multi-label image classification* studies, since it could learn dependency among different labels [65-67]. As shown in **Fig. 2**, a three-layer LSTM was *first* constructed to learn the hierarchical relationships among 6,109 GO families based on the protein encodings pre-trained in ***Module 2***. Each layer of the LSTM had eleven neurons, which were in accordance with those eleven levels in GO families (from LEVEL 1 to LEVEL 11 as shown in **Supplementary Fig. S2**), and the arrows that were provided in the LSTM section of **Fig. 2** indicated that the value of the former neurons (starting point of an arrow) was applied to adjust that of the latter ones (end of that arrow) [68-70]. *Finally*, an ensemble learning approach was applied to integrate sequence similarity into functional prediction, and all proteins could be annotated into a total of 6,109 families.

#### 3.2.2. A Variety of Model Parameters and Their Optimization

Various deep learning strategies were integrated into the development of *AnnoPRO* in this study, which included the *convolutional neural network* (CNN), *deep neural network* (DNN), and *long short-term memory* (LSTM). *First*, CNN contained two *convolution* layers (with their kernel size set to 3 × 3 and stride set to 1) and another two *max-pooling* layers (with their pool length set to 2 and stride set to 1). *Second*, the number of *fully-connected layers* (FC) for developing the DNN models of this study was set to 5. *Third*, the number of layers for constructing the LSTM models of this study was set to 3, and a total of 11 neurons were adopted for each layer.

During model development, a variety of parameters were optimized and systematically provided in **Supplementary Table S3**. *First*, 80% of 92,120 proteins from the CAFA4 benchmark dataset were selected as the training dataset, and the remaining 20% proteins were used as the validation data, which was in accordance with previous study [71]. *Then*, the ‘*mini batch size*’ and ‘*learning rate*’ for ***Module 2*** (Multimodal Protein Encoding Using Multichannel Pre-training) in **Fig. 2** were set to 32 and 0.0002, respectively, with the activation function for both CNN and FC set to *rectified linear unit* (ReLU). *Third*, ‘*mini batch size*’ and ‘*learning rate*’ of ***Module 3*** (Functional Annotation Based on Protein Decoding) in **Fig. 2** were also set to 32 and 0.0002, respectively, with the activation function for LSTM set to *hyperbolic tangent function* (Tanh) [72]. At the *end* of each training epoch, the models’ convergences on validation dataset were carefully monitored, and the model of the best performance was identified using *early stopping* [73]. *Finally*, the *focal loss* was implemented into training process to control the direction of model optimization [74].

#### 3.2.3. The Measurements Facilitating Performance Evaluation

Two well-established measures were adopted in this study for evaluating the model performances, which were widely applied in *critical assessment of functional annotation* (CAFA) challenge [21]. These measures included *area under the precision-recall curve* (AUPRC) & *protein centric maximum F-measure* (F_max_). AUPRC is frequently applied for the evaluation of binary classifiers, especially for assessing the classes of unbalanced data, which is a numeric value between 0 and 1 [75]. The closer the AUPRC is to 1, the better the prediction performance is [75]. F_max_’s strength lies in its interpretability, which is also a numeric value between 0 and 1 [21]. The closer the F_max_ is to 1, the better the prediction performance is. These two measures (AUPRC and F_max_) provided an overall performance assessment of protein functional prediction among different methods, but they were not intuitively enough for predicting a specific protein [76]. Thus, additional measures were adopted into this analysis, which included ‘*recall*’ and ‘*precision*’ [77]. Particularly, ‘*recall*’ evaluated at what percentage the true functions of a protein were successfully predicted, and the closer the recall is to 100%, the more the actual protein functions are annotated. Precision showed what percentage the predicted functions of a protein were true, and the closer precision is to 100%, the more accurately the protein functional annotations are annotated.

## Acknowledgements

This work was supported by National Natural Science Foundation of China (U1909208); Natural Science Foundation of Zhejiang Province (LR21H300001); Leading Talent of the ‘Ten Thousand Plan’ - National High-Level Talents Special Support Plan of China; Fundamental Research Fund for central university (2018QNA7023); Key R&D Program of Zhejiang Province (2020C03010); Double Top-Class University (181201*194232101); Westlake Laboratory (Westlake Laboratory of Life Sciences and Biomedicine); Alibaba-Zhejiang University Joint Research Center of Future Digital Healthcare; Alibaba Cloud; Information Technology Center of Zhejiang University.

## Author Contributions

F.Z. conceived the idea and designed the study; L.Z. proposed *AnnoPRO*’s algorithm; L.Z. and S.S. evaluated the performances of *AnnoPRO*; L.Z., S.S. and M.L. collected related datasets and provided related biological support; L.Z., S.S. and F.Z. wrote the manuscript; P.F., Z.L., S.Z. and Z.Z. built the website. All authors reviewed and approved the manuscript.

## Conflicts of Interest

The authors declare no competing interests.

## Data Availability

The source code for protein functional annotations using *AnnoPRO* is now available on GitHub (https://github.com/idrblab/AnnoPRO), and *AnnoPRO* has also been developed to be an *out-of-the-box* pypi package (https://pypi.org/project/annopro/0.1rc2/). Moreover, an online web server (https://idrblab.org/annopro/) was also developed to enable direct access by all users.

## Notes

https://github.com/idrblab/AnnoPRO

### Competing Interest Statement

The authors have declared no competing interest.

## References

1. Torres M, Yang H, Romero AE, Paccanaro A: **Protein function prediction for newly sequenced organisms**. Nat Mach Intell 2021, 3:1050–1060.

2. Gligorijević V, Renfrew PD, Kosciolek T, Leman JK, Berenberg D, Vatanen T, Chandler C, Taylor BC, Fisk IM, Vlamakis H, et al: **Structure-based protein function prediction using graph convolutional networks**. Nat Commun 2021, 12:3168.

3. Denny P, Feuermann M, Hill DP, Lovering RC, Plun-Favreau H, Roncaglia P: **Exploring autophagy with gene ontology**. Autophagy 2018, 14:419–436.

4. UniProt C: **UniProt: the universal protein knowledgebase in 2021**. Nucleic Acids Res 2021, 49:D480-D489.

5. Lin JS, Lai EM**: Protein-protein interactions: co-immunoprecipitation**. Methods Mol Biol 2017, 1615:211–219.

6. Cui H, Wang Q, Lei Z, Feng M, Zhao Z, Wang Y, Wei G: **DTL promotes cancer progression by PDCD4 ubiquitin-dependent degradation**. J Exp Clin Cancer Res 2019, 38:350.

7. Zhu KY, Palli SR**: Mechanisms, applications, and challenges of insect RNA interference**. Annu Rev Entomol 2020, 65:293–311.

8. You R, Yao S, Xiong Y, Huang X, Sun F, Mamitsuka H, Zhu S: **NetGO: improving large-scale protein function prediction with massive network information**. Nucleic Acids Res 2019, 47:W379–W387.

9. Pearson WR**: Protein function prediction: problems and pitfalls**. Curr Protoc Bioinformatics 2015, 51:4.12.11–14.12.18.

10. Barot M, Gligorijević V, Cho K, Bonneau R**: NetQuilt: deep multispecies network-based protein function prediction using homology-informed network similarity**. Bioinformatics 2021, 37:2414–2422.

11. Kulmanov M, Zhapa-Camacho F, Hoehndorf R**: DeepGOWeb: fast and accurate protein function prediction on the semantic web**. Nucleic Acids Res 2021, 49:W140–W146.

12. Sillitoe I, Furnham N: **FunTree: advances in a resource for exploring and contextualising protein function evolution**. Nucleic Acids Res 2016, 44:D317–D323.

13. Piovesan D, Giollo M, Leonardi E, Ferrari C, Tosatto SC: **INGA: protein function prediction combining interaction networks, domain assignments and sequence similarity**. Nucleic Acids Res 2015, 43:W134–W140.

14. Giri SJ, Dutta P, Halani P, Saha S**: MultiPredGO: deep multi-modal protein function prediction by amalgamating protein structure, sequence, and interaction information**. IEEE J Biomed Health Inform 2021, 25:1832–1838.

15. Kulmanov M, Khan MA, Hoehndorf R, Wren J: **DeepGO: predicting protein functions from sequence and interactions using a deep ontology-aware classifier**. Bioinformatics 2018, 34:660–668.

16. Kulmanov M, Hoehndorf R**: DeepGOPlus: improved protein function prediction from sequence**. Bioinformatics 2020, 36:422–429.

17. Xia W, Zheng L, Fang J, Li F, Zhou Y, Zeng Z, Zhang B, Li Z, Li H, Zhu F**: PFmulDL: a novel strategy enabling multi-class and multi-label protein function annotation by integrating diverse deep learning methods**. Comput Biol Med 2022, 145:105465.

18. Yao S, You R, Wang S, Xiong Y, Huang X, Zhu S: **NetGO 2.0: improving large-scale protein function prediction with massive sequence, text, domain, family and network information**. Nucleic Acids Res 2021, 49:W469–W475.

19. Gene-Ontology C: **The Gene ontology resource: 20 years and still GOing strong**. Nucleic Acids Res 2019, 47:D330–D338.

20. Cui J, Liu S, Tian Z, Zhong Z, Jia J: **ResLT: residual learning for long-tailed recognition**. IEEE Trans Pattern Anal Mach Intell 2022, 2022:3174892.

21. Zhou N, Jiang Y, Bergquist TR, Lee AJ, Kacsoh BZ, Crocker AW, Lewis KA, Georghiou G, Nguyen HN, Hamid MN, et al: **The CAFA challenge reports improved protein function prediction and new functional annotations for hundreds of genes through experimental screens**. Genome Biol 2019, 20:244.

22. Marchler-Bauer A, Bo Y, Han L, He J, Lanczycki CJ, Lu S, Chitsaz F, Derbyshire MK, Geer RC, Gonzales NR, et al: **CDD/SPARCLE: functional classification of proteins via subfamily domain architectures**. Nucleic Acids Res 2017, 45:D200–D203.

23. Lai B, Xu J**: Accurate protein function prediction via graph attention networks with predicted structure information**. Brief Bioinform 2022, 23:bbab502.

24. Zhang C, Freddolino PL, Zhang Y: **COFACTOR: improved protein function prediction by combining structure, sequence and protein-protein interaction information**. Nucleic Acids Res 2017, 45:W291–W299.

25. Hong J, Luo Y, Zhang Y, Ying J, Xue W, Xie T, Tao L, Zhu F**: Protein functional annotation of simultaneously improved stability, accuracy and false discovery rate achieved by a sequence-based deep learning**. Brief Bioinform 2020, 21:1437–1447.

26. Hong J, Luo Y, Mou M, Fu J, Zhang Y, Xue W, Xie T, Tao L, Lou Y, Zhu F**: Convolutional neural network-based annotation of bacterial type IV secretion system effectors with enhanced accuracy and reduced false discovery**. Brief Bioinform 2020, 21:1825–1836.

27. Yu CY, Li XX, Yang H, Li YH, Xue WW, Chen YZ, Tao L, Zhu F**: Assessing the performances of protein function prediction algorithms from the perspectives of identification accuracy and false discovery rate**. Int J Mol Sci 2018, 19:183.

28. Gong Q, Ning W, Tian W**: GoFDR: a sequence alignment based method for predicting protein functions**. Methods 2016, 93:3–14.

29. Tung CC, Kuo SC, Yang CL, Yu JH, Huang CE, Liou PC, Sun YH, Shuai P, Su JC, Ku C, Lin YJ: **Single-cell transcriptomics unveils xylem cell development and evolution**. Genome Biol 2023, 24:3.

30. Du L, Liu Q, Fan Z, Tang J, Zhang X, Price M, Yue B, Zhao K**: Pyfastx: a robust python package for fast random access to sequences from plain and gzipped FASTA/Q files**. Brief Bioinform 2021, 22:bbaa368.

31. Seligmann H**: Alignment-based and alignment-free methods converge with experimental data on amino acids coded by stop codons at split between nuclear and mitochondrial genetic codes**. Biosystems 2018, 167:33-46.

32. Zielezinski A, Vinga S, Almeida J, Karlowski WM: **Alignment-free sequence comparison: benefits, applications, and tools**. Genome Biol 2017, 18:186.

33. Li YH, Li XX, Hong JJ, Wang YX, Fu JB, Yang H, Yu CY, Li FC, Hu J, Xue WW, et al: **Clinical trials, progression-speed differentiating features and swiftness rule of the innovative targets of first-in-class drugs**. Brief Bioinform 2020, 21:649–662.

34. Basharat Z, Akhtar U, Khan K, Alotaibi G, Jalal K, Abbas MN, Hayat A, Ahmad D, Hassan SS**: Differential analysis of Orientia tsutsugamushi genomes for therapeutic target identification and possible intervention through natural product inhibitor screening**. Comput Biol Med 2022, 141:105165.

35. Begum K, Mohl JE, Ayivor F, Perez EE, Leung MY**: GPCR-PEnDB: a database of protein sequences and derived features to facilitate prediction and classification of G protein-coupled receptors**. Database 2020, 2020:baaa087.

36. Mishra S, Rastogi YP, Jabin S, Kaur P, Amir M, Khatun S**: A deep learning ensemble for function prediction of hypothetical proteins from pathogenic bacterial species**. Comput Biol Chem 2019, 83:107147.

37. Wan C, Cozzetto D, Fa R, Jones DT**: Using deep maxout neural networks to improve the accuracy of function prediction from protein interaction networks**. PLoS One 2019, 14:e0209958.

38. Ieremie I, Ewing RM, Niranjan M**: TransformerGO: predicting protein-protein interactions by modelling the attention between sets of gene ontology terms**. Bioinformatics 2022, 38:2269–2277.

39. Sureyya Rifaioglu A, Dogan T, Jesus Martin M, Cetin-Atalay R, Atalay V**: DEEPred: automated protein function prediction with multi-task feed-forward deep neural networks**. Sci Rep 2019, 9:7344.

40. Yao S, You R, Wang S, Xiong Y, Huang X, Zhu S: **NetGO 3.0: protein language model improves large-scale functional annotations**. bioRxiv 2022, **2022**:2022.2012.2005.519073.

41. Bileschi ML, Belanger D, Bryant DH, Sanderson T, Carter B, Sculley D, Bateman A, DePristo MA, Colwell LJ: **Using deep learning to annotate the protein universe**. Nat Biotechnol 2022, 40:932–937.

42. Reher R, Aron AT, Fajtova P, Stincone P, Wagner B, Perez-Lorente AI, Liu C, Shalom IYB, Bittremieux W, Wang M, et al: **Native metabolomics identifies the rivulariapeptolide family of protease inhibitors**. Nat Commun 2022, 13:4619.

43. Hoarfrost A, Aptekmann A, Farfanuk G, Bromberg Y**: Deep learning of a bacterial and archaeal universal language of life enables transfer learning and illuminates microbial dark matter**. Nat Commun 2022, 13:2606.

44. Wang J, Yang Y, Mao JH, Huang ZH, Huang C, Xu W: **CNN-RNN: a unified framework for multi-label image classification**. IEEE Conf Comput Vis Pattern Recognit 2016, 2016:2285–2294.

45. Cao Y, Shen Y**: TALE: transformer-based protein function annotation with joint sequence-Label embedding**. Bioinformatics 2021, 37:2825–2833.

46. Jumper J, Evans R, Pritzel A, Green T, Figurnov M, Ronneberger O, Tunyasuvunakool K, Bates R, Zidek A, Potapenko A, et al: **Highly accurate protein structure prediction with AlphaFold**. Nature 2021, 596:583–589.

47. Kulmanov M, Hoehndorf R**: DeepGOZero: improving protein function prediction from sequence and zero-shot learning based on ontology axioms**. Bioinformatics 2022, 38:i238–i245.

48. Chowdhury R, Bouatta N, Biswas S, Floristean C, Kharkar A, Roy K, Rochereau C, Ahdritz G, Zhang J, Church GM, et al: **Single-sequence protein structure prediction using a language model and deep learning**. Nat Biotechnol 2022, 40:1617–1623.

49. Ba Q, Hei Y, Dighe A, Li W, Maziarz J, Pak I, Wang S, Wagner GP, Liu Y: **Proteotype coevolution and quantitative diversity across 11 mammalian species**. Sci Adv 2022, 8:eabn0756.

50. Kwon D, Lee D, Kim J, Lee J, Sim M, Kim J: **INTERSPIA: a web application for exploring the dynamics of protein-protein interactions among multiple species**. Nucleic Acids Res 2018, 46:W89–W94.

51. Gonzalez JM, Hernandez L, Manzano I, Pedros-Alio C**: Functional annotation of orthologs in metagenomes: a case study of genes for the transformation of oceanic dimethylsulfoniopropionate**. ISME J 2019, 13:1183–1197.

52. Loewenstein Y, Raimondo D, Redfern O, Watson J, Frishman D, Linial M, Orengo C, Thornton J, Tramontano A: **Protein function annotation by homology-based inference**. Genome Biol 2009, 10:207.

53. Schafer MJ, LeBrasseur NK: **The influence of GDF11 on brain fate and function**. GeroScience 2019, 41:1–11.

54. Sinha M, Jang YC, Oh J, Khong D, Wu EY, Manohar R, Miller C, Regalado SG, Loffredo FS, Pancoast JR, et al: **Restoring systemic GDF11 levels reverses age-related dysfunction in mouse skeletal muscle**. Science 2014, 344:649–652.

55. Cash JN, Angerman EB, Kattamuri C, Nolan K, Zhao H, Sidis Y, Keutmann HT, Thompson TB: **Structure of myostatin·follistatin-like 3: N-terminal domains of follistatin-type molecules exhibit alternate modes of binding**. J Biol Chem 2012, 287:1043–1053.

56. Padyana AK, Vaidialingam B, Hayes DB, Gupta P, Franti M, Farrow NA: **Crystal structure of human GDF11**. Acta Crystallogr F Struct Biol Commun 2016, 72:160–164.

57. Cash JN, Rejon CA, McPherron AC, Bernard DJ, Thompson TB**: The structure of myostatin:follistatin 288: insights into receptor utilization and heparin binding**. EMBO J 2009, 28:2662–2676.

58. Suh J, Lee YS**: Similar sequences but dissimilar biological functions of GDF11 and myostatin**. Exp Mol Med 2020, 52:1673–1693.

59. Yun CW, Kim HJ, Lim JH, Lee SH**: Heat shock proteins: agents of cancer development and therapeutic targets in anti-cancer therapy**. Cells 2019, 9:60.

60. Rao HB, Zhu F, Yang GB, Li ZR, Chen YZ: **Update of PROFEAT: a web server for computing structural and physicochemical features of proteins and peptides from amino acid sequence**. Nucleic Acids Res 2011, 39:W385–W390.

61. Bajusz D, Rácz A, Héberger K**: Why is tanimoto index an appropriate choice for fingerprint-based similarity calculations?** J Cheminform 2015, 7:20.

62. Jones W, Chawdhary A, King A**: Optimising the volgenant–jonker algorithm for approximating graph edit distance**. Pattern Recognit Lett 2017, 87:47–54.

63. Dai Z, Cai B, Lin Y, Chen J: **Unsupervised pre-training for detection transformers**. IEEE Trans Pattern Anal Mach Intell 2022, 2022:3216514.

64. Zhang J, Li S**: Air quality index forecast in Beijing based on CNN-LSTM multi-model**. Chemosphere 2022, 308:136180.

65. Kollias D, Zafeiriou S: **Exploiting multi-CNN features in CNN-RNN based dimensional emotion recognition on the OMG in-the-wild dataset**. IEEE Trans Affect Comput 2021, 12:595-606.

66. Xu Y, Hosny A, Zeleznik R, Parmar C, Coroller T, Franco I, Mak RH: **Deep learning predicts lung cancer treatment response from serial medical imaging**. Clin Cancer Res 2019, 25:3266–3275.

67. You Y, Lu C, Wang W, Tang CK**: Relative CNN-RNN: learning relative atmospheric visibility from images**. IEEE Trans Image Process 2019, 28:45–55.

68. Geravanchizadeh M, Roushan H**: Dynamic selective auditory attention detection using RNN and reinforcement learning**. Sci Rep 2021, 11:15497.

69. Gao R, Zhao S, Aishanjiang K, Cai H, Wei T, Zhang Y, Liu Z, Zhou J, Han B, Wang J, et al: **Deep learning for differential diagnosis of malignant hepatic tumors based on multi-phase contrast-enhanced CT and clinical data**. J Hematol Oncol 2021, 14:154.

70. Tsukiyama S, Hasan MM, Fujii S, Kurata H: **LSTM-PHV: prediction of human-virus protein-protein interactions by LSTM with word2vec**. Brief Bioinform 2021, 22:bbab228.

71. Shin HC, Roth HR, Gao M, Lu L, Xu Z, Nogues I, Yao J, Mollura D, Summers RM: **Deep convolutional neural networks for computer-aided detection: CNN architectures, dataset characteristics and transfer learning**. IEEE Trans Med Imaging 2016, 35:1285-1298.

72. De-Ryck T, Lanthaler S, Mishra S: **On the approximation of functions by tanh neural networks**. Neural Netw 2021, 143:732–750.

73. Zhang T, Zhu T, Gao K, Zhou W, Yu PS**: Balancing learning model privacy, fairness, and accuracy with early stopping criteria**. IEEE Trans Neural Netw Learn Syst 2021, 2021:3129592.

74. Lin TY, Goyal P, Girshick R, He K, Dollar P: **Focal loss for dense object detection**. IEEE Trans Pattern Anal Mach Intell 2020, 42:318–327.

75. Ozenne B, Subtil F, Maucort-Boulch D**: The precision-recall curve overcame the optimism of the receiver operating characteristic curve in rare diseases**. J Clin Epidemiol 2015, 68:855–859.

76. Necci M, Piovesan D, Caid P, DisProt C, Tosatto SCE: **Critical assessment of protein intrinsic disorder prediction**. Nat Methods 2021, 18:472–481.

77. Yang H, Chen L, Cheng Z, Yang M, Wang J, Lin C, Wang Y, Huang L, Chen Y, Peng S, et al: **Deep learning-based six-type classifier for lung cancer and mimics from histopathological whole slide images: a retrospective study**. BMC Med 2021, 19:80.

