## Supplementary for "AnnoPRO: an Innovative Strategy for Protein Function Annotation based on Image-like Protein Representation and Multimodal Deep Learning"

*AnnoPRO* was implemented with the libraries of *Keras* and *TensorFlow* as a framework[75]. To accelerate the training process, NVIDIA Tesla P100 PCIe GPUs were used. The total cost of time for model construction was over three months, which is very time-consuming process. However, once the model was constructed, it only takes about one minute to annotate the functions of 1,000 proteins. The source code for protein functional annotations using *AnnoPRO* is now available on GitHub (https://github.com/idrblab/AnnoPRO), and *AnnoPRO* has also been developed to be an *out-of-the-box* pipy package (https://pypi.org/project/annopro/0.1rc2/). Moreover, an online web server (https://idrblab.org/annopro/) was also developed to enable direct access by all users.

### Competing Interests

The authors declare no competing interests.

### References


1. Torres, M., Yang, H., Romero, A.E. & Paccanaro, A. Protein function prediction for newly sequenced organisms. *Nat Mach Intell* **3**, 1050-1060 (2021).
2. Gligorijević, V. et al. Structure-based protein function prediction using graph convolutional networks. *Nat Commun* **12**, 3168 (2021).
3. Denny, P. et al. Exploring autophagy with gene ontology. *Autophagy* **14**, 419-436 (2018).
4. UniProt, C. UniProt: the universal protein knowledgebase in 2021. *Nucleic Acids Res* **49**, D480-D489 (2021).
5. Lin, J.S. & Lai, E.M. Protein-protein interactions: co-immunoprecipitation. *Methods Mol Biol* **1615**, 211-219 (2017).
6. Cui, H. et al. DTL promotes cancer progression by PDCD4 ubiquitin-dependent degradation. *J Exp Clin Cancer Res* **38**, 350 (2019).
7. Zhu, K.Y. & Palli, S.R. Mechanisms, applications, and challenges of insect RNA interference. *Annu Rev Entomol* **65**, 293-311 (2020).
8. You, R. et al. NetGO: improving large-scale protein function prediction with massive network information. *Nucleic Acids Res* **47**, W379-W387 (2019).
9. Pearson, W.R. Protein function prediction: problems and pitfalls. *Curr Protoc Bioinformatics* **51**, 4.12.1-4.12.8 (2015).
10. Barot, M., Gligorijević, V., Cho, K. & Bonneau, R. NetQuilt: deep multispecies network-based protein function prediction using homology-informed network similarity. *Bioinformatics* **37**, 2414-2422 (2021).
11. Kulmanov, M., Zhapa-Camacho, F. & Hoehndorf, R. DeepGOWeb: fast and accurate protein function prediction on the semantic web. *Nucleic Acids Res* **49**, W140-W146 (2021).
12. Sillitoe, I. & Furnham, N. FunTree: advances in a resource for exploring and contextualising protein function evolution. *Nucleic Acids Res* **44**, D317-D323 (2016).
13. Piovesan, D., Giollo, M., Leonardi, E., Ferrari, C. & Tosatto, S.C. INGA: protein function prediction combining interaction networks, domain assignments and sequence similarity. *Nucleic Acids Res* **43**, W134-W140 (2015).
14. Giri, S.J., Dutta, P., Halani, P. & Saha, S. MultiPredGO: deep multi-modal protein function prediction by amalgamating protein structure, sequence, and interaction information. *IEEE J Biomed Health Inform* **25**, 1832-1838 (2021).
15. Kulmanov, M., Khan, M.A., Hoehndorf, R. & Wren, J. DeepGO: predicting protein functions from sequence and interactions using a deep ontology-aware classifier. *Bioinformatics* **34**, 660-668 (2018).
16. Kulmanov, M. & Hoehndorf, R. DeepGOPlus: improved protein function prediction from sequence. *Bioinformatics* **36**, 422-429 (2020).
17. Xia, W. et al. PFmulDL: a novel strategy enabling multi-class and multi-label protein function annotation by integrating diverse deep learning methods. *Comput Biol Med* **145**, 105465 (2022).
18. Yao, S. et al. NetGO 2.0: improving large-scale protein function prediction with massive sequence, text, domain, family and network information. *Nucleic Acids Res* **49**, W469-W475 (2021).
19. Gene-Ontology, C. The Gene ontology resource: 20 years and still GOing strong. *Nucleic Acids Res* **47**, D330-D338 (2019).

20. Cui, J., Liu, S., Tian, Z., Zhong, Z. & Jia, J. ResLT: residual learning for long-tailed recognition. *IEEE Trans Pattern Anal Mach Intell* **2022**, 3174892 (2022).
21. Zhou, N. et al. The CAFA challenge reports improved protein function prediction and new functional annotations for hundreds of genes through experimental screens. *Genome Biol* **20**, 244 (2019).
22. Marchler-Bauer, A. et al. CDD/SPARCLE: functional classification of proteins via subfamily domain architectures. *Nucleic Acids Res* **45**, D200-D203 (2017).
23. Lai, B. & Xu, J. Accurate protein function prediction via graph attention networks with predicted structure information. *Brief Bioinform* **23**, bbab502 (2022).
24. Zhang, C., Freddolino, P.L. & Zhang, Y. COFACTOR: improved protein function prediction by combining structure, sequence and protein-protein interaction information. *Nucleic Acids Res* **45**, W291-W299 (2017).
25. Hong, J. et al. Protein functional annotation of simultaneously improved stability, accuracy and false discovery rate achieved by a sequence-based deep learning. *Brief Bioinform* **21**, 1437-1447 (2020).
26. Hong, J. et al. Convolutional neural network-based annotation of bacterial type IV secretion system effectors with enhanced accuracy and reduced false discovery. *Brief Bioinform* **21**, 1825-1836 (2020).
27. Yu, C.Y. et al. Assessing the performances of protein function prediction algorithms from the perspectives of identification accuracy and false discovery rate. *Int J Mol Sci* **19**, 183 (2018).
28. Gong, Q., Ning, W. & Tian, W. GoFDR: a sequence alignment based method for predicting protein functions. *Methods* **93**, 3-14 (2016).
29. Tung, C.C. et al. Single-cell transcriptomics unveils xylem cell development and evolution. *Genome Biol* **24**, 3 (2023).
30. Du, L. et al. Pyfastx: a robust python package for fast random access to sequences from plain and gzipped FASTA/Q files. *Brief Bioinform* **22**, bbaa368 (2021).
31. Seligmann, H. Alignment-based and alignment-free methods converge with experimental data on amino acids coded by stop codons at split between nuclear and mitochondrial genetic codes. *Biosystems* **167**, 33-46 (2018).
32. Zielezinski, A., Vinga, S., Almeida, J. & Karlowski, W.M. Alignment-free sequence comparison: benefits, applications, and tools. *Genome Biol* **18**, 186 (2017).
33. Li, Y.H. et al. Clinical trials, progression-speed differentiating features and swiftness rule of the innovative targets of first-in-class drugs. *Brief Bioinform* **21**, 649-662 (2020).
34. Basharat, Z. et al. Differential analysis of Orientia tsutsugamushi genomes for therapeutic target identification and possible intervention through natural product inhibitor screening. *Comput Biol Med* **141**, 105165 (2022).
35. Begum, K., Mohl, J.E., Ayivor, F., Perez, E.E. & Leung, M.Y. GPCR-PEnDB: a database of protein sequences and derived features to facilitate prediction and classification of G protein-coupled receptors. *Database* **2020**, baaa087 (2020).
36. Mishra, S. et al. A deep learning ensemble for function prediction of hypothetical proteins from pathogenic bacterial species. *Comput Biol Chem* **83**, 107147 (2019).
37. Wan, C., Cozzetto, D., Fa, R. & Jones, D.T. Using deep maxout neural networks to improve the accuracy of function prediction from protein interaction networks. *PLoS One* **14**, e0209958 (2019).
38. Ieremie, I., Ewing, R.M. & Niranjan, M. TransformerGO: predicting protein-protein interactions by modelling the attention between sets of gene ontology terms.

*Bioinformatics* **38**, 2269-2277 (2022).
39. Sureyya Rifaioglu, A., Dogan, T., Jesus Martin, M., Cetin-Atalay, R. & Atalay, V. DEEPred: automated protein function prediction with multi-task feed-forward deep neural networks. *Sci Rep* **9**, 7344 (2019).
40. Yao, S. et al. NetGO 3.0: protein language model improves large-scale functional annotations. *bioRxiv* **2022**, 2022.12.05.519073 (2022).
41. Bileschi, M.L. et al. Using deep learning to annotate the protein universe. *Nat Biotechnol* **40**, 932-937 (2022).
42. Reher, R. et al. Native metabolomics identifies the rivulariapeptolide family of protease inhibitors. *Nat Commun* **13**, 4619 (2022).
43. Hoarfrost, A., Aptekmann, A., Farfanuk, G. & Bromberg, Y. Deep learning of a bacterial and archaeal universal language of life enables transfer learning and illuminates microbial dark matter. *Nat Commun* **13**, 2606 (2022).
44. Wang, J. et al. CNN-RNN: a unified framework for multi-label image classification. *IEEE Conf Comput Vis Pattern Recognit* **2016**, 2285-2294 (2016).
45. Cao, Y. & Shen, Y. TALE: transformer-based protein function annotation with joint sequence-Label embedding. *Bioinformatics* **37**, 2825-2833 (2021).
46. Jumper, J. et al. Highly accurate protein structure prediction with AlphaFold. *Nature* **596**, 583-589 (2021).
47. Kulmanov, M. & Hoehndorf, R. DeepGOZero: improving protein function prediction from sequence and zero-shot learning based on ontology axioms. *Bioinformatics* **38**, i238-i245 (2022).
48. Chowdhury, R. et al. Single-sequence protein structure prediction using a language model and deep learning. *Nat Biotechnol* **40**, 1617-1623 (2022).
49. Ba, Q. et al. Proteotype coevolution and quantitative diversity across 11 mammalian species. *Sci Adv* **8**, eabn0756 (2022).
50. Kwon, D. et al. INTERSPIA: a web application for exploring the dynamics of protein-protein interactions among multiple species. *Nucleic Acids Res* **46**, W89-W94 (2018).
51. Gonzalez, J.M., Hernandez, L., Manzano, I. & Pedros-Alio, C. Functional annotation of orthologs in metagenomes: a case study of genes for the transformation of oceanic dimethylsulfoniopropionate. *ISME J* **13**, 1183-1197 (2019).
52. Loewenstein, Y. et al. Protein function annotation by homology-based inference. *Genome Biol* **10**, 207 (2009).
53. Schafer, M.J. & LeBrasseur, N.K. The influence of GDF11 on brain fate and function. *GeroScience* **41**, 1-11 (2019).
54. Sinha, M. et al. Restoring systemic GDF11 levels reverses age-related dysfunction in mouse skeletal muscle. *Science* **344**, 649-652 (2014).
55. Cash, J.N. et al. Structure of myostatin·follistatin-like 3: N-terminal domains of follistatin-type molecules exhibit alternate modes of binding. *J Biol Chem* **287**, 1043-1053 (2012).
56. Padyana, A.K. et al. Crystal structure of human GDF11. *Acta Crystallogr F Struct Biol Commun* **72**, 160-164 (2016).
57. Cash, J.N., Rejon, C.A., McPherron, A.C., Bernard, D.J. & Thompson, T.B. The structure of myostatin:follistatin 288: insights into receptor utilization and heparin binding. *EMBO J* **28**, 2662-2676 (2009).
58. Suh, J. & Lee, Y.S. Similar sequences but dissimilar biological functions of GDF11 and myostatin. *Exp Mol Med* **52**, 1673-1693 (2020).

59. Yun, C.W., Kim, H.J., Lim, J.H. & Lee, S.H. Heat shock proteins: agents of cancer development and therapeutic targets in anti-cancer therapy. *Cells* **9**, 60 (2019).

60. Rao, H.B., Zhu, F., Yang, G.B., Li, Z.R. & Chen, Y.Z. Update of PROFEAT: a web server for computing structural and physicochemical features of proteins and peptides from amino acid sequence. *Nucleic Acids Res* **39**, W385-W390 (2011).

61. Bajusz, D., Rácz, A. & Héberger, K. Why is tanimoto index an appropriate choice for fingerprint-based similarity calculations? *J Cheminform* **7**, 20 (2015).

62. Jones, W., Chawdhary, A. & King, A. Optimising the volgenant–jonker algorithm for approximating graph edit distance. *Pattern Recognit Lett* **87**, 47-54 (2017).

63. Dai, Z., Cai, B., Lin, Y. & Chen, J. Unsupervised pre-training for detection transformers. *IEEE Trans Pattern Anal Mach Intell* **2022**, 3216514 (2022).

64. Zhang, J. & Li, S. Air quality index forecast in Beijing based on CNN-LSTM multi-model. *Chemosphere* **308**, 136180 (2022).

65. Kollias, D. & Zafeiriou, S. Exploiting multi-CNN features in CNN-RNN based dimensional emotion recognition on the OMG in-the-wild dataset. *IEEE Trans Affect Comput* **12**, 595-606 (2021).

66. Xu, Y. et al. Deep learning predicts lung cancer treatment response from serial medical imaging. *Clin Cancer Res* **25**, 3266-3275 (2019).

67. You, Y., Lu, C., Wang, W. & Tang, C.K. Relative CNN-RNN: learning relative atmospheric visibility from images. *IEEE Trans Image Process* **28**, 45-55 (2019).

68. Geravanchizadeh, M. & Roushan, H. Dynamic selective auditory attention detection using RNN and reinforcement learning. *Sci Rep* **11**, 15497 (2021).

69. Gao, R. et al. Deep learning for differential diagnosis of malignant hepatic tumors based on multi-phase contrast-enhanced CT and clinical data. *J Hematol Oncol* **14**, 154 (2021).

70. Tsukiyama, S., Hasan, M.M., Fujii, S. & Kurata, H. LSTM-PHV: prediction of human-virus protein-protein interactions by LSTM with word2vec. *Brief Bioinform* **22**, bbab228 (2021).

71. Shin, H.C. et al. Deep convolutional neural networks for computer-aided detection: CNN architectures, dataset characteristics and transfer learning. *IEEE Trans Med Imaging* **35**, 1285-1298 (2016).

72. De-Ryck, T., Lanthaler, S. & Mishra, S. On the approximation of functions by tanh neural networks. *Neural Netw* **143**, 732-750 (2021).

73. Zhang, T., Zhu, T., Gao, K., Zhou, W. & Yu, P.S. Balancing learning model privacy, fairness, and accuracy with early stopping criteria. *IEEE Trans Neural Netw Learn Syst* **2021**, 3129592 (2021).

74. Lin, T.Y., Goyal, P., Girshick, R., He, K. & Dollar, P. Focal loss for dense object detection. *IEEE Trans Pattern Anal Mach Intell* **42**, 318-327 (2020).

75. Tang, H. et al. TensorFlow solver for quantum PageRank in large-scale networks. *Sci Bull* **66**, 120-126 (2021).

76. Ozenne, B., Subtil, F. & Maucort-Boulch, D. The precision-recall curve overcame the optimism of the receiver operating characteristic curve in rare diseases. *J Clin Epidemiol* **68**, 855-859 (2015).

77. Necci, M., Piovesan, D., Caid, P., DisProt, C. & Tosatto, S.C.E. Critical assessment of protein intrinsic disorder prediction. *Nat Methods* **18**, 472-481 (2021).

78. Yang, H. et al. Deep learning-based six-type classifier for lung cancer and mimics from histopathological whole slide images: a retrospective study. *BMC Med* **19**, 80 (2021).

**Table 1**. A comparison among the performances of *AnnoPRO* and eight available methods/tools. The values indicating the best performances among all methods/tools were highlighted in BOLD, and *AnnoPRO* performed consistently the best in all Gene Ontology (GO) classes (BP, CC, MF) under both evaluating criteria ($F_{max}$, AUPRC). All methods/tools were ordered according to their publication dates. BP: *biological process*; CC: *cellular component*; MF: *molecular function*; $F_{max}$: *protein centric maximum F-measure*; AUPRC: *area under the precision-recall curve*. Those tools marked by an asterisk (*) indicated that their source-codes for model construction were not fully provided, which made it impossible for us to train models on experimental functional annotations that appeared before Oct 22, 2019, and their performances (evaluated by $F_{max}$ and AUPRC) were assessed by directly uploading those experimental function annotations asserted between Oct 22, 2019 and May 31, 2022 to the online server of those annotation tools. Among those eight existing methods/tools, the best performing ones under each category were highlighted by *underline*.

| Method/Tool | Date of Publication | BP | | CC | | MF | |
|---|---|---|---|---|---|---|---|
| | | $F_{max}$ | AUPRC | $F_{max}$ | AUPRC | $F_{max}$ | AUPRC |
| *DiamondBLAST* | Nov, 2014 | 0.549 | 0.183 | 0.550 | 0.186 | 0.729 | 0.112 |
| *DeepGO* | Feb, 2018 | 0.362 | 0.213 | 0.501 | 0.434 | 0.384 | 0.325 |
| *DeepGOCNN* | Jan, 2020 | 0.369 | 0.294 | 0.516 | 0.460 | 0.382 | 0.362 |
| *DeepGOPlus* | Jan, 2020 | 0.593 | 0.561 | 0.588 | 0.502 | 0.628 | 0.627 |
| *TALE* | Mar, 2021 | 0.391 | 0.307 | 0.562 | 0.587 | 0.472 | 0.458 |
| *NetGO2** | Jul, 2021 | 0.497 | 0.434 | 0.574 | 0.508 | 0.667 | 0.674 |
| *PFmulDL* | Mar, 2022 | 0.324 | 0.257 | 0.590 | 0.608 | 0.412 | 0.371 |
| *NetGO3** | Dec, 2022 | 0.540 | 0.500 | 0.579 | 0.535 | 0.687 | 0.726 |
| *AnnoPRO* | This Study | **0.609** | **0.574** | **0.746** | **0.749** | **0.763** | **0.755** |

**Table 2.** A comparison among the performances of *AnnoPRO* and two *state-of-the-art* methods (*DeepGOPlus* and *PFmulDL*) on predicting two groups of '*Independent Testing*' data (**SameSP** and **DiffSP**). **SameSP** had 1,859 proteins from 17 species covered by '*Training*' and '*Validation*' datasets of this study; **DiffSP** included 3,764 proteins from the remaining 997 species unique in '*Independent Testing*' data of this study. Those values indicating the best performance among all three methods were highlighted in BOLD, and *AnnoPRO* performed the best in the vast majority of the Gene Ontology (GO) classes (BP, CC & MF) under both evaluating criteria ($F_{max}$, AUPRC). BP: *biological process*; CC: *cellular component*; MF: *molecular function*.

| | Method | BP | | CC | | MF | |
|---|---|---|---|---|---|---|---|
| | | **$F_{max}$** | **AUPRC** | **$F_{max}$** | **AUPRC** | **$F_{max}$** | **AUPRC** |
| **SameSP** | *DeepGOPlus* | **0.612** | **0.593** | 0.539 | 0.470 | 0.668 | 0.698 |
| | *PFmulDL* | 0.347 | 0.286 | 0.573 | 0.603 | 0.436 | 0.402 |
| | *AnnoPRO* | 0.610 | 0.589 | **0.759** | **0.772** | **0.835** | **0.829** |
| **DiffSP** | *DeepGOPlus* | 0.538 | 0.469 | 0.684 | 0.622 | 0.517 | 0.429 |
| | *PFmulDL* | 0.261 | 0.176 | 0.593 | 0.580 | 0.354 | 0.273 |
| | *AnnoPRO* | **0.602** | **0.552** | **0.742** | **0.741** | **0.749** | **0.739** |

**Table 3**. The prediction of the '*heparin binding*' function (GO:0008201) for the wild type GDF8 (***GDF8-WT***) and its two GDF8 mutants (***GDF8-Mutant-1*** & ***GDF8-Mutant-2***) using *AnnoPRO* and three representative methods. 'Success' denotes that the gain/loss-of-function is successfully predicted by the corresponding method, while 'Fail' indicates that it is incorrectly predicted. As demonstrated, significant functional variations among ***GDF8-WT***, ***GDF8-Mutant-1***, and ***GDF8-Mutant-2*** can only be "successfully" captured by our newly developed AnnoPRO.

| **Methods** | ***GDF8-WT**** | ***GDF8-Mutant-1**** | ***GDF8-Mutant-2**** |
|---|---|---|---|
| *DeepGOPlus* | Fail | Fail | Success |
| *PFmulDL* | Fail | Fail | Success |
| *NetGO3* | Success | Success | Fail |
| ***AnnoPRO*** | Success | Success | Success |

* The wild type GDF8 (***GDF8-WT***) is a growth differentiation factor of 375 amino acids. There are two GDF8 mutants (***GDF8-Mutant-1*** & ***GDF8-Mutant-2***). ***GDF8-Mutant-1*** contains eight mutations (D267N, F268L, T277S, E312Q, H328Q, G355D, E357Q, and A366G) which locate far away from the binding interface between GDF8 and follistatin-288 (FS288). The interaction between ***GDF8-WT*** and FS288 forms a protein complex to further bind to heparin. This is the molecular mechanism underlying ***GDF8-WT***'s key GO term: '*heparin binding*' (GO:0008201). Because all eight mutations were far away from the binding interface between GDF8 and FS288, it is expected that the '*heparin binding*' function remains in ***GDF8-Mutant-1***[58]. Meanwhile, ***GDF8-Mutant-2*** contains three mutations (F315Y, V316M, and L318M, on the binding surface between GDF8 and FS288) which are reported as the key residues indicating protein's '*heparin binding*' function[58]. In other words, it is expected that ***GDF8-Mutant-2*** loses its wild type's '*heparin binding*' function[58]. All in all, there is gain-of-function of '*heparin binding*' in both ***GDF8-WT*** and ***GDF8-Mutant-1***, while there is loss-of-function in ***GDF8-Mutant-2***.

**Table 4**. A comparison among the predictive performances of *AnnoPRO* and three representative methods for the functional annotations of two well-known *growth differentiation factors* (GDF8, GDF11). Those values indicating the best performances among all methods were highlighted in BOLD, and *AnnoPRO* performed the best in the vast-majority (11/12) of the Gene Ontology (GO) classes (BP, CC, MF) under both evaluating criteria (recall, precision). All methods were ordered based on their publication dates. BP: *biological process*; CC: *cellular component*; MF: *molecular function*; GDF8: *growth differentiation factor 8*; GDF11: *growth differentiation factor 11*.

| **Protein Name** | **Methods** | **BP** | | **CC** | | **MF** | |
|---|---|---|---|---|---|---|---|
| | | **Recall** | **Precision** | **Recall** | **Precision** | **Recall** | **Precision** |
| GDF8 | *DeepGOPlus* | 0.578 | 0.320 | 0.333 | **1.000** | 0.389 | 0.333 |
| | *PFmulDL* | 0.333 | 0.198 | 0.667 | 0.400 | 0.444 | 0.444 |
| | *NetGO3* | 0.351 | 0.806 | **1.000** | 0.375 | **1.000** | 0.783 |
| | ***AnnoPRO*** | **0.898** | **0.898** | **1.000** | 0.731 | **1.000** | **1.000** |
| GDF11 | *DeepGOPlus* | 0.402 | 0.306 | 0.625 | 0.714 | 0.222 | **1.000** |
| | *PFmulDL* | 0.404 | 0.494 | 0.875 | 0.412 | 0.556 | 0.833 |
| | *NetGO3* | 0.553 | 0.547 | 0.750 | 0.750 | 0.778 | **1.000** |
| | ***AnnoPRO*** | **0.621** | **0.952** | **1.000** | **0.833** | **1.000** | **1.000** |

**Table 5**. A comparison among the predictive performances of *AnnoPRO* and three representative methods for the functional annotations of two well-known *heat shock 70kDa proteins* (HSPA1A, HSPA2). Those values indicating the best performances among all methods were highlighted in BOLD, and *AnnoPRO* performed the best in the vast-majority (9/12) of the Gene Ontology (GO) classes (BP, CC, MF) under both evaluating criteria (recall, precision). All methods were ordered based on their publication dates. BP: *biological process*; CC: *cellular component*; MF: *molecular function*; HSPA1A: *heat shock 70 kDa protein 1A*; HSPA2: *heat shock 70 kDa protein 2*.

| **Protein Name** | **Methods** | **BP** | | **CC** | | **MF** | |
|---|---|---|---|---|---|---|---|
| | | **Recall** | **Precision** | **Recall** | **Precision** | **Recall** | **Precision** |
| HSPA1A | *DeepGOPlus* | 0.358 | 0.357 | 0.410 | 0.889 | 0.605 | 0.812 |
| | *PFmulDL* | 0.635 | 0.457 | 0.615 | 0.800 | 0.814 | 0.500 |
| | *NetGO3* | 0.286 | **0.876** | **0.634** | 0.605 | 0.809 | 0.884 |
| | ***AnnoPRO*** | **0.641** | 0.715 | 0.595 | **0.962** | **0.917** | **0.936** |
| HSPA2 | *DeepGOPlus* | 0.375 | 0.284 | 0.394 | **0.867** | 0.765 | 0.765 |
| | *PFmulDL* | 0.344 | 0.386 | 0.419 | 0.812 | 0.788 | 0.605 |
| | *NetGO3* | 0.346 | 0.605 | 0.419 | 0.684 | 0.757 | 0.903 |
| | ***AnnoPRO*** | **0.470** | **0.851** | **0.594** | 0.670 | **0.868** | **0.943** |

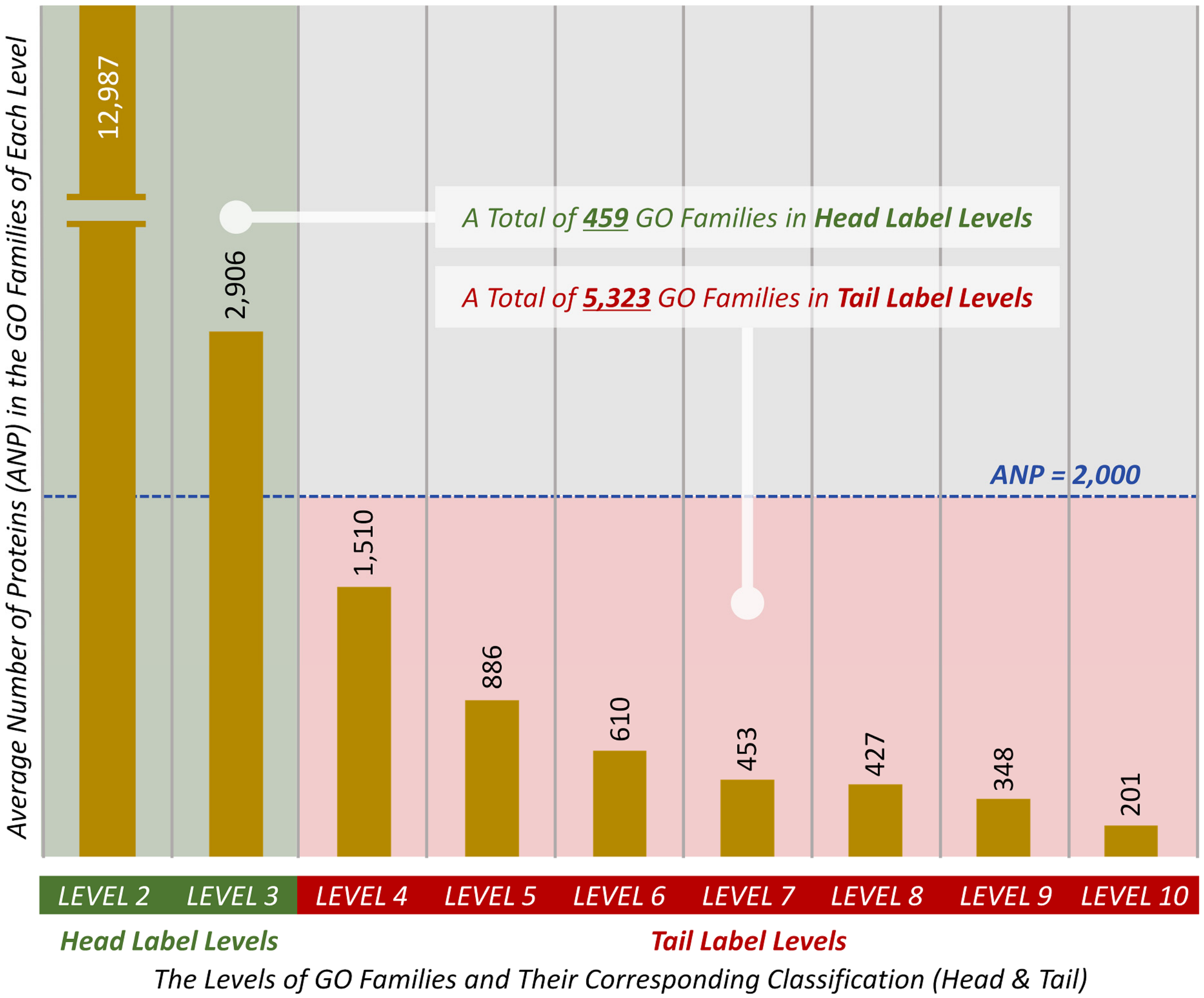


**Figure 1**. Average numbers of proteins (ANP) in the GO families of nine different levels (LEVEL 2 to LEVEL 10 as shown in **Supplementary Figure S2**). There was a clear descending trend of ANPs from the top level (LEVEL 2) to the bottom one (LEVEL 10). Since the ANP of one family indicated its representativeness among all families, this figure denoted a gradual decrease of the representativeness of a family with the penetration into deeper level. Therefore, these nine levels could be classified into two groups based on their ANPs: the "*Head Label Levels*" (ANP of their GO families ≥ 2,000) and the "*Tail Label Levels*" (ANP of their GO families < 2,000). As shown, the total number (5,323) of GO families in the "*Tail Label Levels*" was >10 times larger than that (459) of the "*Head Label Levels*", and such kind of data distribution induced a serious '*long-tail problem*' as described in the previous pioneering publication[20].

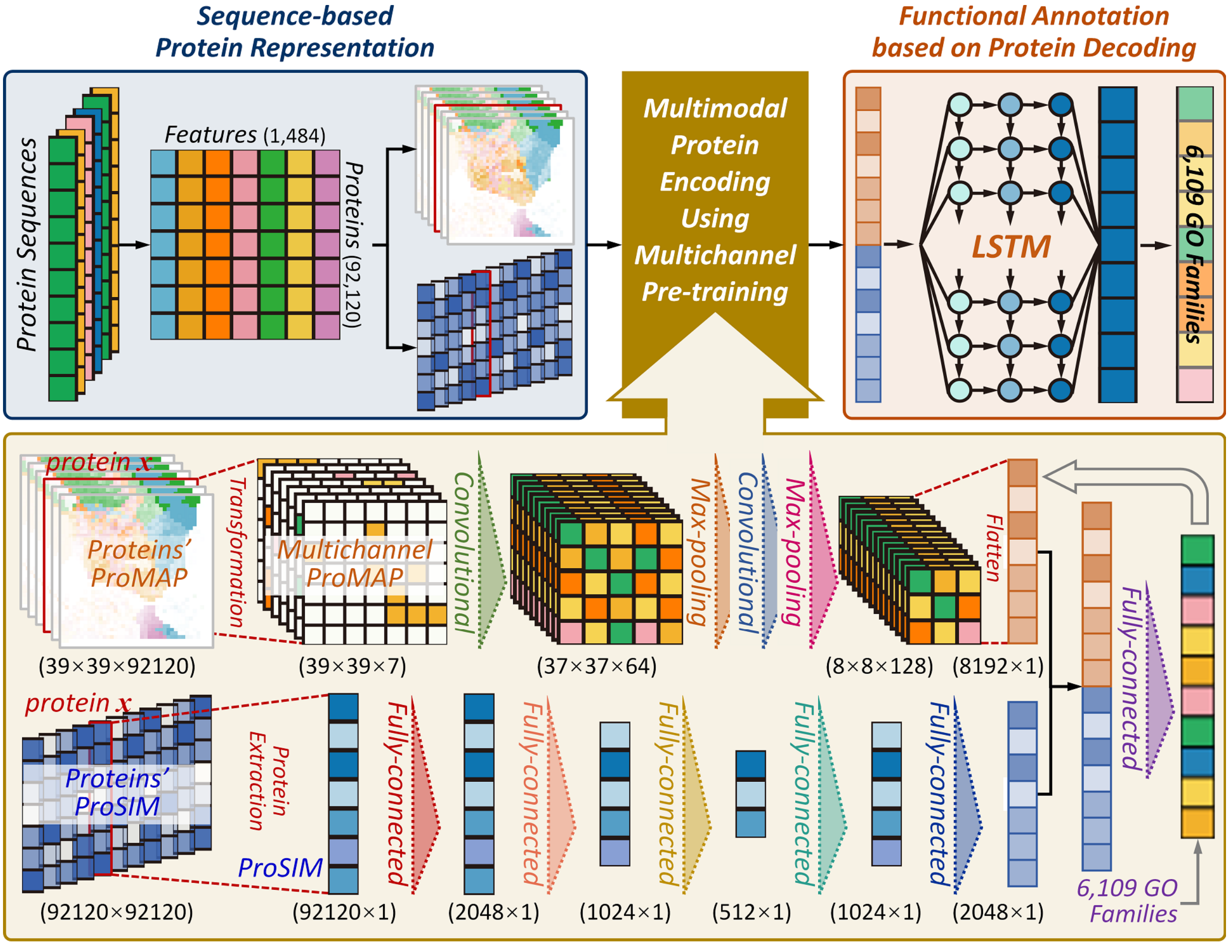


**Figure 2**. The multimodal and multichannel deep learning framework of *AnnoPRO*. There were three consecutive modules (***M1*** to ***M3***). (***M1***) sequence-based protein representation that realized the conversion of all protein sequences to the image-like protein representation. (***M2***) multimodal protein encoding using multichannel pre-training. Using the *ProMAP* and *ProSIM* generated here for proteins, a multimodal and multichannel deep learning architecture was constructed based on a seven-channel *convolutional neural network* (7C-CNN) & a *deep neural network* of five fully-connected layers (5FC-DNN) to pre-train the features of all CAFA4 proteins by integrating their annotation data of GO functional families. (***M3***) functional annotation based on protein decoding. The protein features pre-trained using the dual-path multimodal encoding layer in second module were concatenated and then fed into a *long short-term memory recurrent neural network* (LSTM) to enable a comprehensive multilabel annotation of proteins to 6,109 functional GO families.

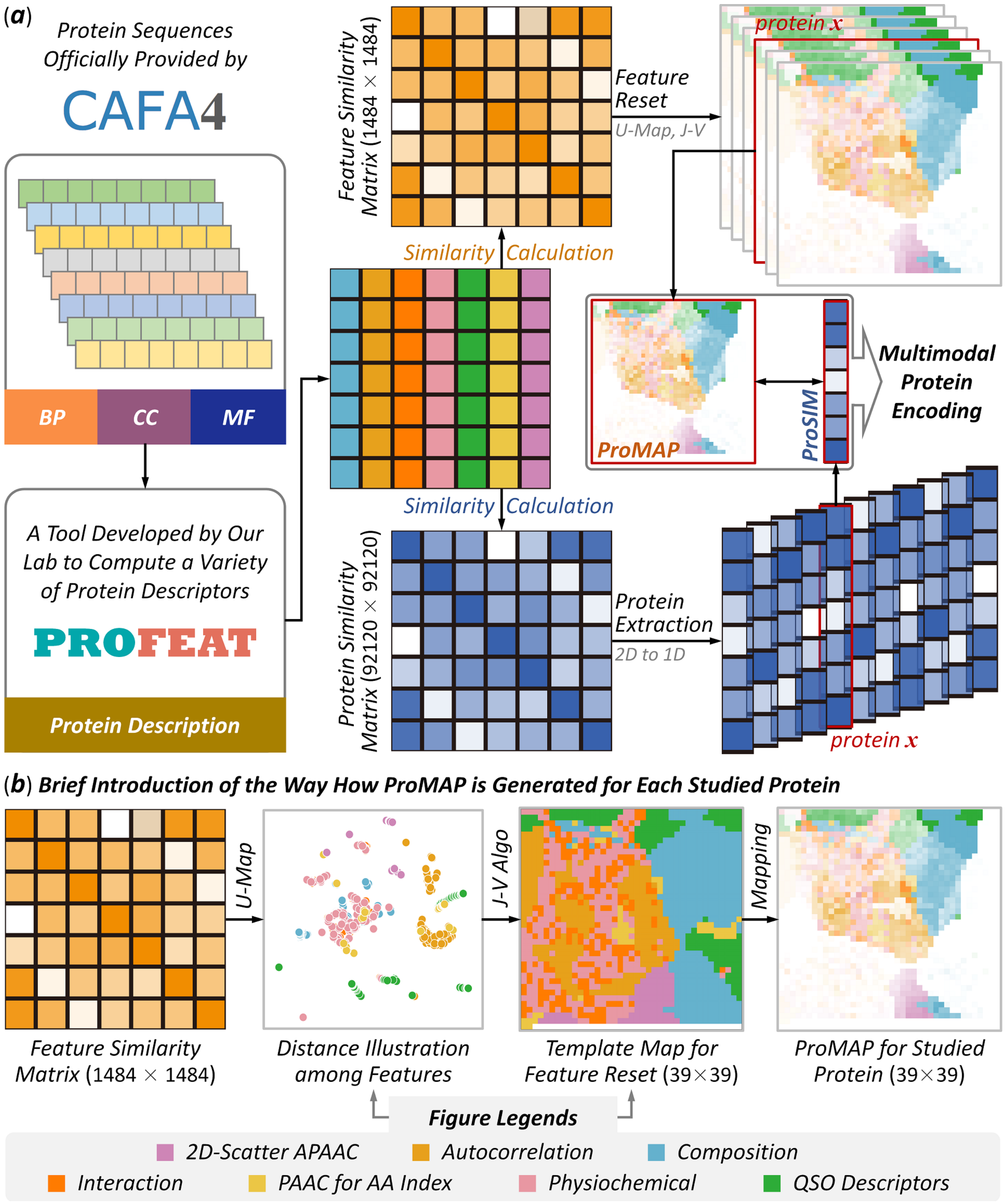


**Figure 3**. A schematic illustration of the procedure used in this study facilitating sequence-based protein representation. (***a***) a concrete description on the novel strategy proposed in this study for representing protein sequences based on calculating the similarities among proteins and features. Particularly, a total of 92,120 protein sequences were *first* collected from the official website of CAFA4, and their descriptors were computed by a popular tool developed by our research lab[60]; two similarity matrices (showing similarity among features & proteins) were *then* generated; the feature similarity matrix was *further* applied to reset the location of all features in a 2D map, and

any sequence could therefore be represented as a 39×39 square image (which was named in this study as '*ProMAP*' for each protein); the protein similarity matrix was *also* adopted to extract a 1D vector, and any protein sequence could be represented as a vector of 92,120 features (which was named here as '*ProSIM*' for each protein); and both *ProMAP* and *ProSIM* were *finally* used to facilitate the subsequent "*multimodal protein encoding*" (the second module shown in **Figure 2**). (***b***) a brief introduction of the way how a '*ProMAP*' was created for each protein. Particularly, feature similarity matrix was *first* applied to illustrate the distances among any two features based on U-Map[61]; a template image/map for feature reset was *then* generated by *Jonker-Volgenant* (*J-V*) algorithm[62] and U-Map; and *ProMAP* was *finally* produced for each protein by mapping the intensities of all features to their corresponding locations in the template image.

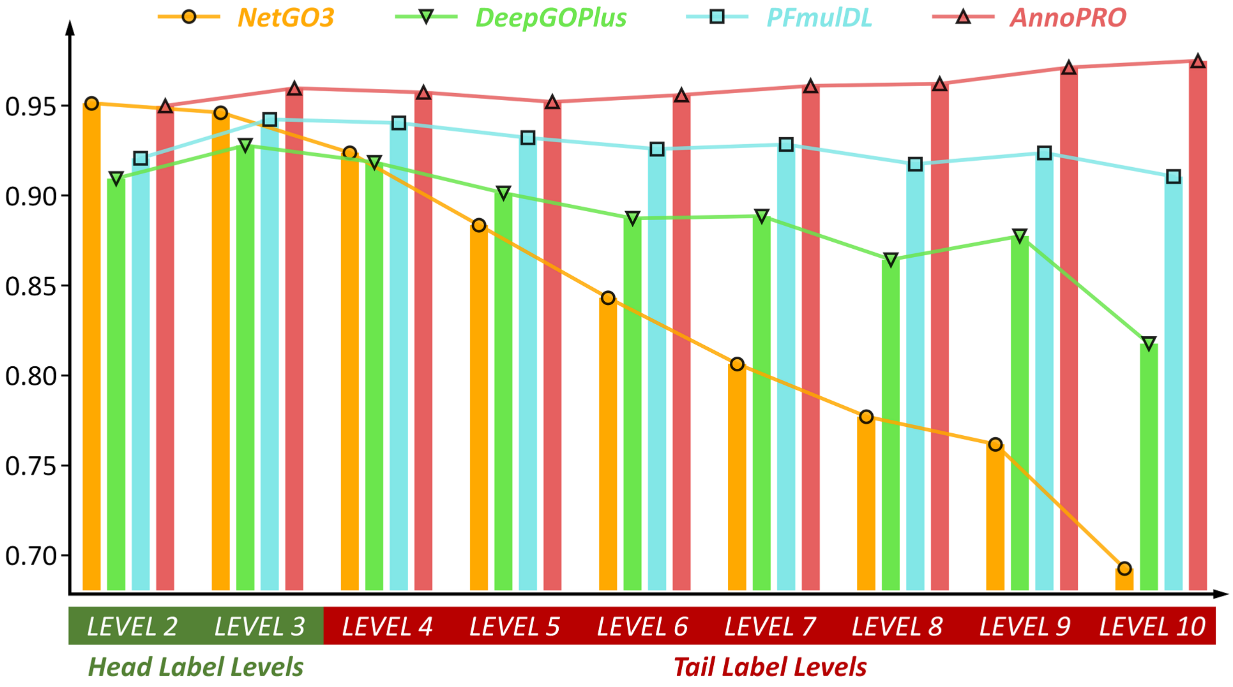


**Figure 4**. A comparison among the performances of *AnnoPRO* and three representative methods. The performances were represented using AUC values in predicting the experimentally validated new protein functions that were not included in CAFA4 data, and the performances of *AnnoPRO*, *DeepGOPlus*, *NetGO3* and *PFmulDL* were highlighted in light red, light green, orange and light blue, respectively. For the GO families in the '*Head Label Levels*' (LEVEL 2 & LEVEL 3 shown in **Supplementary Figure S2**), the performance of *AnnoPRO* was roughly as good as that of the other three methods (1.4~4.1% improvements in most cases, but 0.1% decline in one single case). For the GO families in the '*Tail Label Levels*' (LEVEL 4 to LEVEL 10 shown in **Supplementary Figure S2**), *AnnoPRO* demonstrated the consistently superior performance among four methods (1.7~28.2% improvements in all cases). Particularly, 13 (61.9%) out of all 21 improvements were over 5%, and 6 (28.6%) out of 21 improvements were more than 10%. Therefore, *AnnoPRO* was identified *superior* in significantly improving the annotation performances of the families in '*Tail Label Levels*' without sacrificing that of the '*Head Label Levels*', which was highly expected to make contribution to solving the long-standing '*long-tail problem*'[20] in functional annotation.

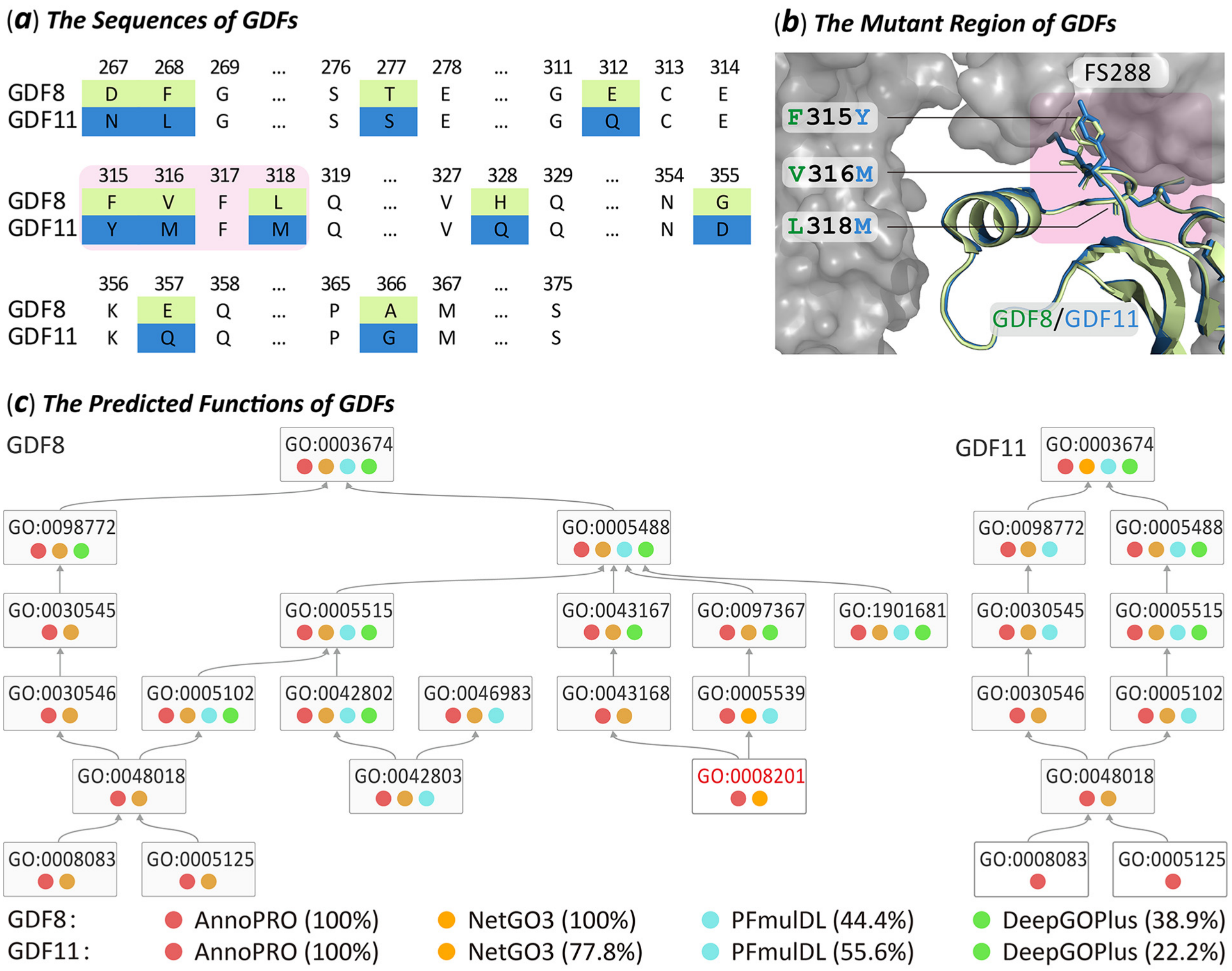


**Figure 5**. Performance assessment of four methods using two well-known *growth differentiation factors* (GDF8, GDF11). As reported, the interaction between GDF8 and follistatin-288 (FS288) formed a protein complex to bind 'heparin', which defined the molecular mechanisms underlying GDF8's key GO family: '*heparin binding*' (GO:0008201)[55]. Different from GDF8, the varied residues in GDF11 made it unable to interact with FS288, and it therefore suffered from the loss of the '*heparin binding*' function[56]. (***a***) sequence alignment between GDF8 and GDF11, where varied residues between two GDFs were marked in light green and blue background, respectively. Three varied residue pairs (F315Y, V316M & L318M on the binding surface between GDF8 and FS288) which were found as key residue indicating GDFs' '*heparin binding*' function[58], were shown in pink background. (***b***) structural superimposition between GDF8 (light green) & GDF11 (blue) and their interactions with FS288 (gray surface). As highlighted in pink background, those three key residue pairs (F315Y, V316M & L318M) located right in the binding interface between GDF and FS288. (***c***) the results of functional annotation predicted by four studied methods. If a GO family is successfully predicted by a method, a colored circle will be adopted to indicate the prediction result. Particularly, a successful prediction made by *AnnoPRO*, *NetGO3*, *PFmulDL* or *DeepGOPlus* was indicated by a circle of light red, orange, light blue or light green, respectively. As shown, *AnnoPRO* is the only one that can successfully predict all GO families for both GDFs.
